# From Syllables to Words: EEG Evidence of Different Age Trajectories in Speech Tracking and Statistical Learning in Infants at High and Low Likelihood for Autism

**DOI:** 10.1101/2025.11.20.689632

**Authors:** Michel Godel, Ana Fló, Lucas Benjamin, Ghislaine Dehaene-Lambertz, Marie Schaer

**Author notes:** these authors have contributed equally to this work (Ghislaine Dehaene-Lambertz, Marie Schaer).

## Abstract

Delayed onset of canonical babbling and first words is often reported in infants later diagnosed with autism spectrum disorder. Identifying the neural mechanisms underlying language acquisition in autism is therefore critical to inform early diagnosis, prognosis, and intervention strategies. In this study, we investigated two speech processing mechanisms previously identified as atypical in children and adults with autism: the neural ability to track syllables; and statistical learning, the capacity to detect speech regularities beneath surface variability. We recorded 83 longitudinal high-density EEGs from 44 infants (2.5–22.6 months) at high (HL) and low (LL) likelihood for autism and assessed their verbal outcomes at 20 months. Neural entrainment was measured at syllable and word frequencies during exposure to a multi-speaker stream of concatenated tri-syllabic words, followed by a word recognition test using ERP recording. Our findings revealed reduced tracking abilities at the syllabic level in HL infants, a measure that correlated with verbal outcomes. While HL infants did not exhibit deficits in statistical learning itself, they displayed reduced novelty orientation during the word recognition test, indicated by a reduced late ERP. By contrast, multi-talker variability temporarily disrupted word segmentation around 12 months in LL infants, but not in HL infants, potentially reflecting decreased sensitivity to human voices variability in the HL group. These results emphasize the importance of longitudinal protocols employing online, implicit measures to track the hierarchical stages of speech processing in both HL and LL infants.

## Introduction

Autism Spectrum Disorder (hereafter: autism) is a neurodevelopmental condition marked by early and pervasive challenges in social interaction and communication, coupled with repetitive behaviors and restricted interests (1). The prevalence of autism has risen over the past decades to an estimated 3.2% (2,3). It is frequently associated with language difficulties that greatly vary across individuals and age (4,5). Atypical babbling and reduced word recognition are among the earliest indicators of autism (6,7). Identifying and addressing these early difficulties is crucial to mitigating their long-term detrimental cascading effects on later verbal and non-verbal abilities (8–10). Achieving this, however, requires a deeper understanding of the neural mechanisms underlying language acquisition in infants who eventually develop autism compared to typically developing peers.

A key step in language acquisition is the ability to discover the deep linguistic structure that lies beneath a variable surface. In particular, a primary challenge is to perceive the chain of discrete words embedded in a continuous speech flow, a difficult problem for autistic individuals vividly described by Donna Williams: “the way my brain had broken down sentences into words left me with a strange and sometimes unintelligible message” (11). This capacity relies both on the correct identification of the successive phonemes in the spoken flow and their attribution to a given word, as, unlike written text, where spaces delineate words, spoken language lacks clear perceptual boundaries. Successful word segmentation relies on a complex and hierarchical integration of prosodic, phonetic, lexical, and contextual cues (12). Among these cues, Saffran and colleagues (1996) highlighted the role of statistical regularities in the speech stream, showing that after just two minutes of exposure to a stream of randomly concatenated four tri-syllabic non-words, 8-month-old infants could distinguish these words from syllable combinations spanning word boundaries (13). This ability was attributed to the learning of transition probabilities between adjacent syllables—the computation of the likelihood of syllable B coming after syllable A in the artificial speech, P(B|A). In their experiment, where each word was followed by one of three other words, transition probabilities were 1.0 within a word and 0.33 across word boundaries. This ability has put forward the contribution of statistical learning to speech segmentation, notably in preverbal infants. It was later shown that this capacity is already available in sleeping neonates (14,15). Nowadays, statistical learning is recognized as a general learning mechanism, available at any age (16) and observed in a wide range of perceptual domains (17–19). It operates largely automatically, as it has been observed even in sleeping and comatose states (15,20). Importantly, a causal role of statistical learning in language acquisition difficulties has been proposed in several studies of developmental disorders (21–24).

Considering this background, infants who eventually develop autism may face challenges at two levels of auditory processing. First, they may struggle at a perceptual stage: following the rhythm of speech and robustly encoding syllables. Second, statistical learning itself might be impaired. Prospective longitudinal studies of infants at high likelihood for autism (HL) have become the standard methodology to explore how neural processing takes place before the emergence of a reliable diagnosis (25,26). HL status usually stems from a family history of autism and/or a condition strongly associated with autism, like some specific genetic syndromes (27).

Regarding the first level, several EEG studies have reported decreased cortical synchronization to speech syllabic modulations in both autistic children and infants at high likelihood for autism (28–31). Specifically, these studies identified a negative relationship between theta-range (4-7 Hz) power in response to naturalistic speech stimuli and verbal abilities in autistic children. This diminished synchronization may stem from an excitatory/inhibitory neuronal imbalance in auditory cortices (32–34), as well as from anomalies in early auditory perception. For instance, studies have observed jittering or distorted frequency encoding in auditory brainstem responses in autism, which could disrupt subsequent processing stages, such as a correct identification of the phonemes (35–37).

A growing body of literature also emphasizes atypical statistical learning in individuals with autism, particularly when using linguistic stimuli (38). After repeated exposures to a syllable stream, 10 y.o. autistic children showed a magneto-encephalographic neural response that was not modulated by the statistical properties of the syllable sequences, indicating a deficit in statistical learning (39). Exposing autistic children and HL infants to similar stimuli, two independent fMRI studies found that the left temporo-parietal cortex, left amygdala, and basal ganglia were less activated by statistical cues (40,41). However, the difference between groups was mainly due to a lack of activation in autistic children and might be related to low-level sensory difficulties classically described in autism, amplified by the scanner noise rather than to a genuine lack of statistical learning capacities, as discussed by the authors themselves. Importantly, most studies exploring either cortical synchronization or statistical learning have focused on autistic children rather than HL infants, i.e., well beyond the age at which those mechanisms support language acquisition.

Building on the evidence linking statistical learning to early language development and its potential disruptions in autism, we sought to investigate the preverbal developmental trajectory of this ability in infants at low and high likelihood of autism. Using high-density EEG, we conducted a prospective longitudinal study to evaluate how infants process an artificial speech stream similar to the one used by Saffran et al.’s (1996), during their first two years of life. Our study had four primary goals: first, to map the infant developmental trajectory of auditory statistical learning, which remains incompletely understood even in typically developing infants (42); second, to identify the specific levels of word learning at which HL infants may show difficulties; third, to determine whether these difficulties were stable, worsened, or improved over time compared to LL infants, and fourth, to assess whether any of these early indicators could predict later verbal difficulties.

We recorded 83 high-density EEG sessions from 44 infants (19 LL, 25 HL) aged 2.5 to 22.6 months, using an experimental paradigm originally designed for sleeping newborns (15,43,44). During the experiment, infants were exposed to a stream composed of randomly concatenated syllables (random stream or RND) and to an artificial speech stream consisting of syllables of constant duration that formed tri-syllabic non-words (structure stream or STR) (**Figure 1A**). Following this learning phase, they listened to isolated triplets corresponding to words from the stream and to part-words, which spanned word boundaries. This design allowed us to obtain several neural measures, which should help identify the specific difficulties faced by autistic children. First, the regular presentation of syllables at a fixed duration elicits increased power and Phase Locking Values (PLV) at the syllable frequency (4 Hz), providing an efficiency measure of neural synchronization with the speech signal. Second, previous research has shown that if the regular word structure is detected, neural entrainment at the word frequency emerges, resulting in increased PLV at the word frequency (4 Hz / 3 syllables = 1.33 Hz). This measure reflects the ability to segment the stream into words through statistical computations (15,16,45,46). Finally, comparing ERPs to isolated words and part-words enables to assess subsequent word recognition through familiarity/surprise responses (15). For each electrophysiological measure, we assessed its association with participants’ receptive and expressive verbal outcomes collected at 18-21 months of age.

**Figure 1.**
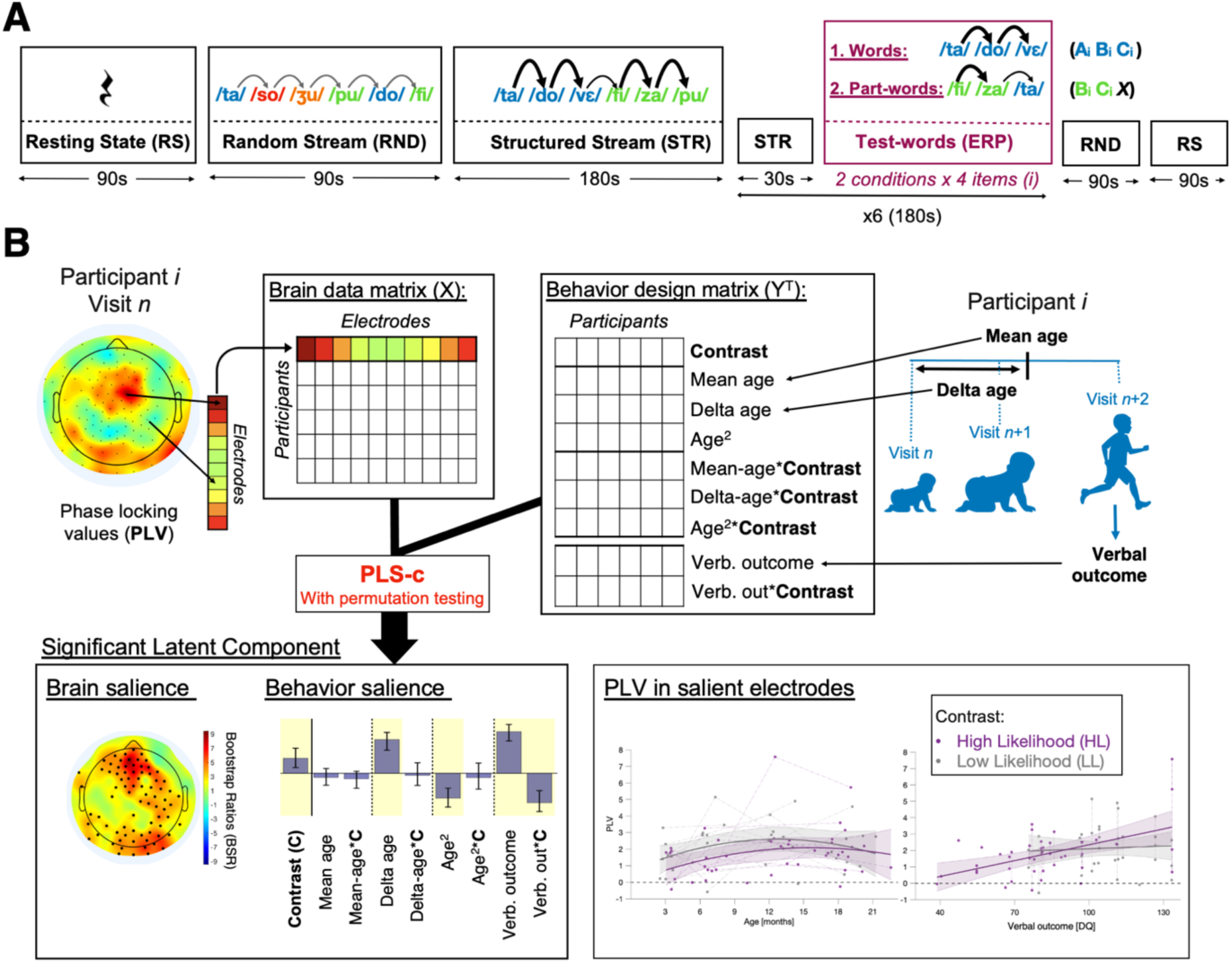
A. Experimental procedure and multivariate statistical analyses. The learning part was sandwiched by a silent resting state (RS) and a random stream (RND) with even transition probabilities between syllables. This design accounted for the potential effect of time during the experiment and changes in vigilance state on neural entrainment measures. The learning segment consisted of a long structured (STR) stream where syllables were organized into four three-syllable words presented in random order with no repetition. Following this, six test-blocks were presented, each comprising 8 triplets from the words and part-words conditions with 2-second silences interleaved between items. To sustain learning, 30-second short STR streams were interspersed between test blocks. A 4.5s fade-in/out at the borders of each stream was included to minimize any perceptual anchor effect. The full procedure lasted ∼17 minutes. Arrows’ width schematically represents the transition probability magnitude. **B. Pipeline for longitudinal partial least square correlation (PLS-c) analysis.** Details are provided in the Methods section.

To increase the cognitive demands of our task and elicit potentially larger differences between LL and HL participants, we introduced random speaker changes across syllables. This acoustic variability was intended to add a layer of complexity that may pose a particular challenge for autistic individuals to filter out (47,48). Despite this variability, LL participants were expected to disregard voice changes, as previous studies have shown that infants can normalize phonetic representations across voices (49), enabling them to focus on the transition probabilities between syllables and extract the statistical structure of the stream. Notably, statistical learning has been demonstrated in neonates under these conditions of voice variation (44). It is worth noting, however, that our study was not designed to isolate or quantify the specific impact of speaker variability on statistical learning, as the experimental design did not include a baseline control condition omitting this acoustic variation.

To account for the multiple variables of the design, data were analyzed using longitudinal partial least square correlations (PLS-c, **Figure 1B**). We included participants’ age, group (HL/LL), and verbal outcome to explain the EEG signal (50,51). Our analyses aim not only to provide a detailed characterization of syllable tracking and statistical learning in autism HL infants compared to typical infants, but also to track the evolution of these fundamental skills over the first two years of life in both groups of infants.

## Results

### Neural entrainment

Analyses of the entrainment at the syllabic rate (4 Hz) and the word rate (1.3 Hz) followed an identical logic. Phase-locking value (PLV) was employed as a measure of neural entrainment, as it directly quantifies neural synchronization to the stimuli. We examined whether neural entrainment was present in the different conditions across all participants while tracing the developmental trajectories of these responses across ages. To achieve this, we used partial least square correlation (PLS-c) (51). Briefly, PLS-c is a data-driven multivariate modelling approach designed to identify significant patterns of electrode clusters (from a brain data matrix containing electrophysiological measures, here PLV) and their associations with “behavioral” variables (from a behavioral design matrix, here age-related parameters). Patterns of brain x behavior associations are called latent components, and their statistical significance is evaluated using permutation testing (n=1000, Bonferroni correction for number of components tested, alpha=.006). Brain and behavioral variables respective contributions to any significant latent component are tested with bootstrapping (500 random samples and replacement), with bootstrap ratios (BSR) greater than 2.3 indicating a stable contribution (for details, see the Materials and Methods section). The analysis aimed to confirm neural entrainment at the syllabic rate during both the random (RND) and structure (STR) streams and, more importantly, at the word rate during the STR stream only. Second, we investigated group differences (LL vs. HL) in entrainment by using a PLS-c with the entrainment at a given frequency as brain data and a behavioral design matrix including the group contrast, the age terms, and the verbal outcome variable. Finally, we assessed the dynamics of statistical learning by tracking changes in neural entrainment throughout the experiment.

### Neural entrainment to syllables

The PLS-c analysis testing for syllabic entrainment in the RND and STR streams recovered one latent component (p < .001, r = .75, 93.1% explained covariance, **Figure 2A-B**). There was a significant contrast effect (entrainment at 4Hz vs. adjacent frequency bins, bootstrap ratio, BSR: 88.1), confirming syllable rate entrainment at the whole sample level. Interestingly, we observed a quadratic age effect (age² term) on the contrast, with a convex trajectory peaking around 12 months (BSR for contrast*age²: -4.0). Separate PLS-C analyses for each stream (i.e., RND and STR) yielded similar results, showing a comparable convex age trajectory in both streams (**Figure 2 -figure supplement 1**). Because some infants were asleep during the recording session, particularly at younger ages, we performed a supplementary control analysis restricted to this sleeping subsample (n = 25 recordings, **Figure 2 -figure supplement 2**). This PLS-c also identified a significant latent component (p<.001, r=.78, 85.1% explained covariance), with a significant contrast effect (BSR: 30.3) and a significant negative contrast*age² interaction (BSR: −2.5). These findings confirm that the convex age trajectory observed in the main analysis remains present and observable even in sleeping infants.

**Figure 2:**
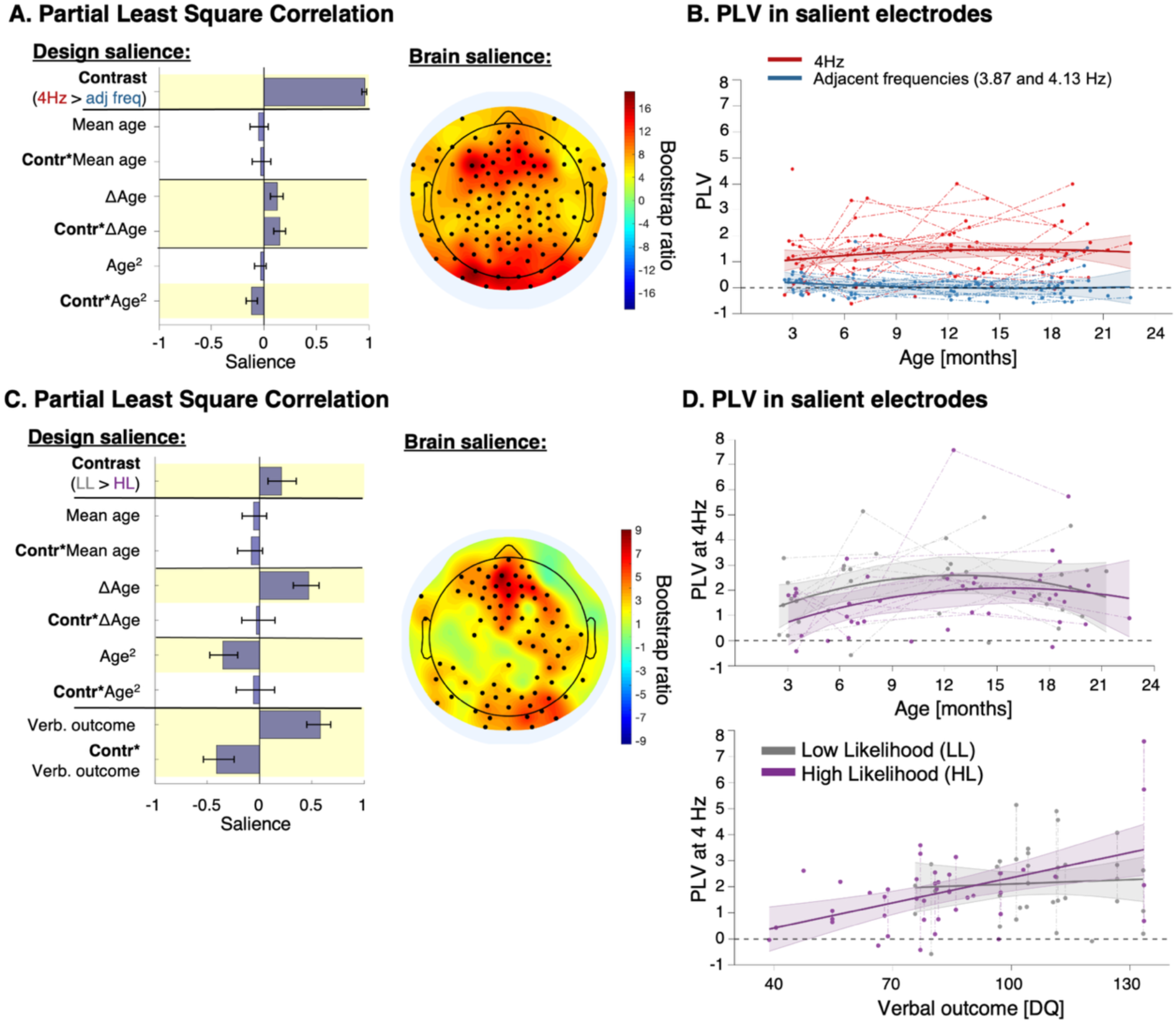
Neural entrainment to syllable rate (4 Hz): Main effect across all participants (Top row); Group differences (bottom row). **A.** Design salience (left) and brain salience (right topography) derived from the significant latent component for neural entrainment to the syllable rate (4 Hz) using the targeted frequency (4 Hz) versus adjacent frequencies as a contrast. Significance (i.e., salience) was established through bootstrapping. Bars represent the mean of 500 random salience samples with replacement bootstrapping, and error bars indicate the 95% confidence interval. Yellow shading highlights variables that significantly contribute to the latent component, defined by a Bootstrap Ratio (BSR; mean of bootstrapping divided by standard deviation) > 2.3. The topography of the BSR values shows electrodes significantly contributing to the latent component (indicated by black dots, BSR > 2.3). **B.** Individual raw Phase Locking Values (PLVs) extracted from the salient electrodes identified by the latent component (black dots in the topography on 2A) are displayed. A fitted curve is included for visualization purposes only, produced by a mixed-effects model with a 95% confidence interval. This curve is intended solely to aid visualization, as the statistical relationships between EEG and behavioral variables are determined by the PLS-C analysis. **C.** PLS-C analysis of the differences between HL and LL groups is presented following the same format as in A. **D.** Individual raw PLVs extracted from the salient electrodes (black dots in subfigure 2C) are shown for visualization purposes only. At every age, a verbal developmental quotient (DQ) of 100 is expected in the general population.

Regarding syllable entrainment differences between groups, we identified one significant latent component (p<.001; r=.50; 61.5% explained covariance, **Figure 2C-D**). This latent component was characterized by a significant effect of group (BSR: 3.1). LL infants showed overall stronger syllable entrainment compared to HL participants. There was no significant age or age*group effect, indicating that syllable entrainment age-trajectories followed a similar convex shape in both groups. Moreover, the latent component comprised a positive effect of verbal outcome (BSR: 10.0), as well as a negative group*verbal outcome interaction effect (BSR: -5.9). This indicates that lower syllable entrainment was associated with poorer verbal performances at 18-21 months of age, predominantly in the HL group. To rule out the possibility that the association between syllable entrainment and verbal outcome was driven by concurrent measures taken at 18–21 months, we re-ran the PLS-c analysis excluding EEG data from the final visit (n=54 recordings kept). The resulting latent component remained significant (p=.001) and showed contributions from behavioral and EEG variables that were highly similar to those observed in the previous analysis, with a verbal outcome BSR of 7.1 and a group*verbal-outcome interaction BSR of −6.3 (**Figure 2 -figure supplement 3**).

### Neural entrainment to words

The PLS-c on 1.33Hz entrainment vs. the entrainment at adjacent frequency bins during the STR stream resulted in one significant latent component (p = .006, r = .33, 30.7% explained covariance). There was a significant contrast effect (BSR: 29.6; **Figure 3A-B**), confirming that at the whole sample level, infants’ brains phased locked to the word rate and thus learned the regularities. This analysis also revealed a quadratic effect of age in the opposite direction of that observed for syllable entrainment, which showed a convex pattern (contrast × age² BSR: – 4.0; **Figure 2A-B**). By contrast, the age-related trajectory for word entrainment was concave, reaching a nadir at 12 months (contrast*age² BSR: 3.6). To confirm the specificity of the word effect, we performed a PLS-c analysis at 1.3 Hz during the RND condition, in which no entrainment was expected. This analysis didn’t reveal any significant latent component. We further investigated word entrainment in sleeping participants (n=25 recordings), which yielded one significant latent component (p<.001, r=.63, 37.3% explained covariance, **Figure 3 -figure supplement 1**). Centro-frontal electrode contributed to this component, with a high contrast BSR (24.9), confirming a similar word entrainment pattern in the sleeping subsample. The contrast contrast*age² was also positive but not significant (1.1), suggesting a trend toward a U-shape age trajectory with a 12-month nadir in sleeping infants. Additional 18-21 month recordings would be required to confirm this trend.

**Figure 3:**
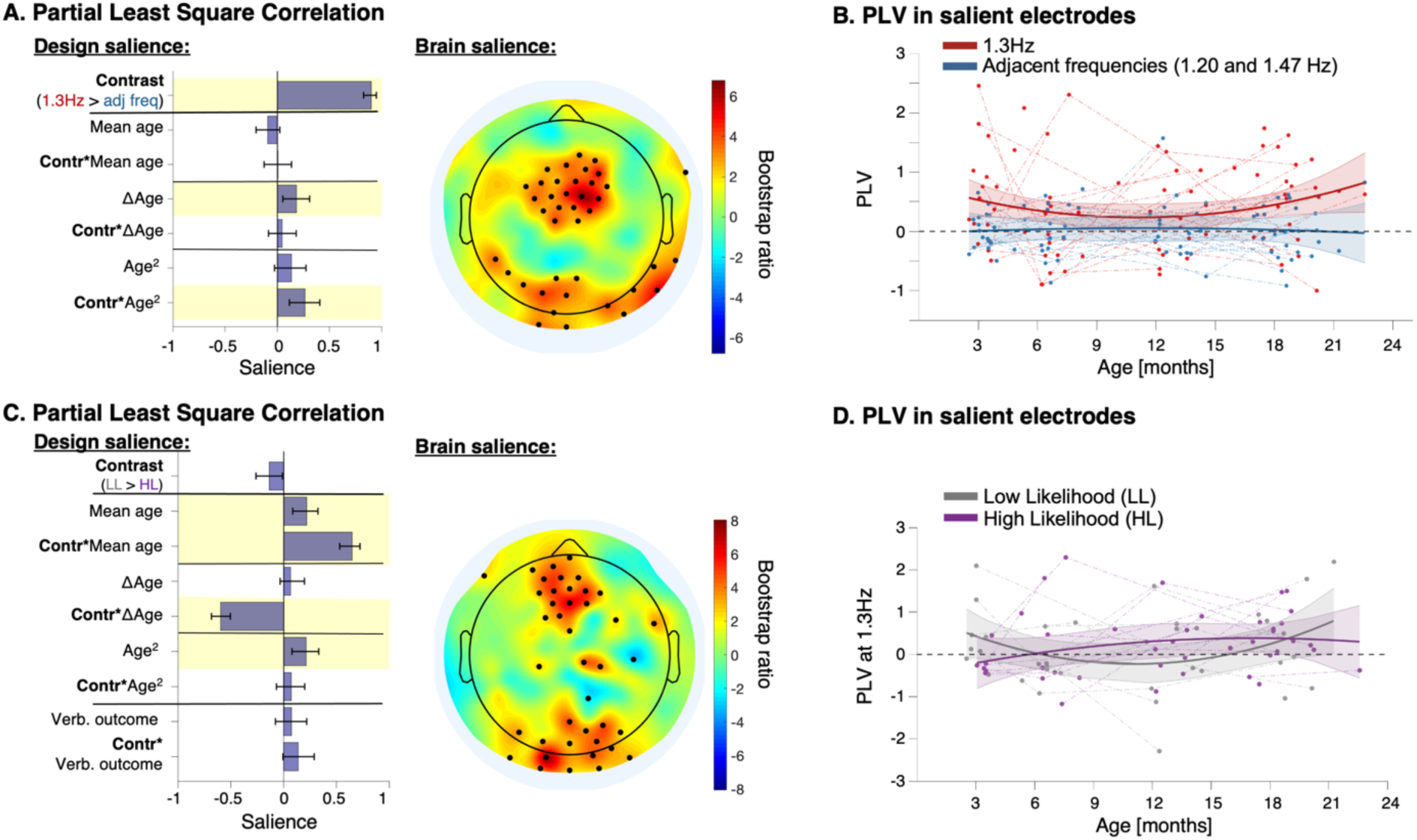
Neural entrainment to word rate (1.3Hz): Main effect across all participants. (**Top row); Group differences (bottom row). A.** Design salience (left) and brain salience (right) derived (through bootstrapping) from the significant latent component for neural entrainment to word rate. **B**. Individual raw PLVs extracted from the salient electrodes given by the latent component (black dots on subfigure A) with a fitted curve, for visualization purposes only. **C.** PLS-c applied on PLV at 1.3Hz with adjacent frequencies subtracted, using group as contrast, and verbal outcome (verbal developmental quotient [DQ] collected at 18-21 months) added as a design variable**. D.** Raw individual data extracted from the salient electrodes of the latent component (black dots on subfigure 2C) with a linear regression curve fitted for illustration purpose only.

Using group as the contrast variable (LL vs HL), we identified a significant latent component (p=.002; r=.50; 27.7% explained covariance) (**Figure 3C-D**), but no reliable group effect (BSR<2.3). This indicates globally homogeneous neural tracking of words across infancy in both groups. There was also no effect of verbal outcome, nor significant interaction verbal outcome *group.

Given previous evidence of impaired statistical learning in autism, we conducted a supplementary PLS-c analysis restricted to the HL group to confirm that HL participants exhibited statistical learning abilities. Using the contrast (1.3Hz vs. adjacent frequencies) and the same age-related design variables, we confirmed significant word entrainment within the HL group (one significant latent component with p=.002, r=.50 and 38.2% explained covariance; contrast BSR=12.9, **Figure 3 -figure supplement 2**). Moreover, the salient electrodes contributing to the latent component showed a spatial distribution closely matching that observed in Figures 3A and 3C, confirming that the core neural mechanisms supporting statistical learning were consistent across groups. However, the significant age*group interaction, along with two additional effects: mean-age*group (BSR: 14.2) capturing cross-sectional age differences, and delta-age*group (BSR: –13.0), suggests distinct developmental trajectories between the two groups. Word entrainment in LL participants followed a more concave (U-shaped) age-trajectory than in HL participants (**Figure 3D**). This interpretation was supported by a supplementary model comparison using Akaike Information Criterion (AIC) to evaluate constant, linear, and quadratic mixed-effects models on raw phase locking values (PLV) extracted from salient electrodes (black dots in **Figure 3C**) within each group (52). For HL participants, the constant model had the lowest AIC (106.4) with a significant intercept above zero (.27 estimate, p=.021). In contrast, for LL participants, the quadratic model was best (AIC =93.4) with significant effects for the intercept (estimate 1.33, p=.025), age (estimate - .24, p=.036) and age^2^ (.01 estimate, p=.049). A PLS-c analysis restricted to the HL group (contrast: 1.33 Hz vs. adjacent frequency bins) further confirmed the absence of a U-shaped trajectory, showing no interaction with age² (BSR: <2.3, **Figure 3 -figure supplement 2**). In summary, statistical learning was reliably present and stable in HL participants, while LL participants showed a transient dip in performance, pointing to divergent developmental trajectories rather than differences in learning capacity.

### Time course of the entrainment along the experiment

As a post-hoc analysis, we explored the temporal dynamics of statistical learning across the session using PLV data at each 1.5-second timeframe within a 120-second time window, averaged across the salient electrodes identified in the previous global analysis. Mixed-effects models fitted at each time frame comparing entrainment at 1.33Hz versus the adjacent frequency bins revealed that word-level entrainment emerged approximately 90 seconds after the onset of the structured stream (Supplementary material, **Figure S3)**.

To gain further insight into the learning dynamics over time in the two groups, we conducted a PLS-c analysis comparing the syllabic and word entrainment time courses between LL and HL participants. HL/LL group was used as a contrast, while the brain data matrix (X) used PLV in both time and space dimensions, i.e., PLV in each electrode at each timeframe. For syllable entrainment, the analysis revealed a significant latent component (*p*<.001; see Supplementary Material **S4**). The group, age, and interaction BSRs were similar to those observed in the previous PLS-c presented in **Figure 2C**. The group, age and verbal outcome parameters were mainly correlated (BSR>2.3) with the neural entrainment occurring ∼90 seconds after the onset of the STR stream, coinciding with the time participants began tracking word boundaries (Supplementary material, **Figure S3**). This result suggests that the group differences in syllable entrainment, as shown in **Figure 2C-D**, as their associations with verbal outcome are modulated by the structure of the stream (STR *versus* RND). We ran one additional PLS-c for each stream separately, using group as contrast. In both streams, the PLS-c yielded a significant latent component (p<.001 for RND and p=.002 for STR), with a positive group effect (BSR>2.3) in both latent components (Supplementary material, **Figure S5**). Most strikingly, the group*verbal outcome parameter reached significance exclusively within the STR latent component (BSR:-7.3). These results suggest that while syllable tracking is generally decreased in HL infants across both streams, its association with verbal outcome is prominently driven by the stream containing words (STR).

In contrast, a PLS-c analysis of the word-level entrainment time course revealed no significant latent components when comparing the two groups.

### ERP analyses

Visual inspection of the grand-average ERP across all recordings showed an early response characterized by a frontal positivity accompanied by a posterior negativity, corresponding to the auditory response and developing over approximately the first second (word duration=750 ms). It was followed by a late component displaying a reversed spatial pattern from 1500 to 3000 ms (see Supplementary Material, **Figure S6**). ERP analyses using either condition or group as contrast variables were conducted in two temporal time windows: an early window sensitive to auditory processing [0-1000ms], and a later window [1500-3000ms] typically associated with higher-order cognitive responses, such as novelty detection and surprise-related activity (53).

### Early ERP

We first tested whether a significant difference between word and part-word conditions was observed across all participants in the early [0-1000ms] time window (grand average topographies are displayed in Supplementary materials, **Figure S6**). We performed a longitudinal PLS-c contrasting part-word and word conditions. This analysis identified one significant latent component (p<.001; R=.54; 63.0% explained covariance, **Figure 4**). However, this component did not show any significant contribution from the condition contrast or its interaction with age variables (BSR<2.3). Instead, only the age-related variables (BSRs for age: -12.6; delta-age: -29.5; age²: 11.1) significantly contributed to the latent component. This indicates an age-related decline in the amplitude of both the frontal positivity and the posterior negativity, following a slightly quadratic concave shape (**Figure 4B**). There was no evidence for any modulatory effect of the part-word/word condition on this developmental pattern. The same PLS-c, conducted separately in sleeping (n=25) and awake subsamples (n=58), yielded no significant latent component, indicating a consistent absence of early response to word novelty in both sleeping and awake infants.

**Figure 4.**
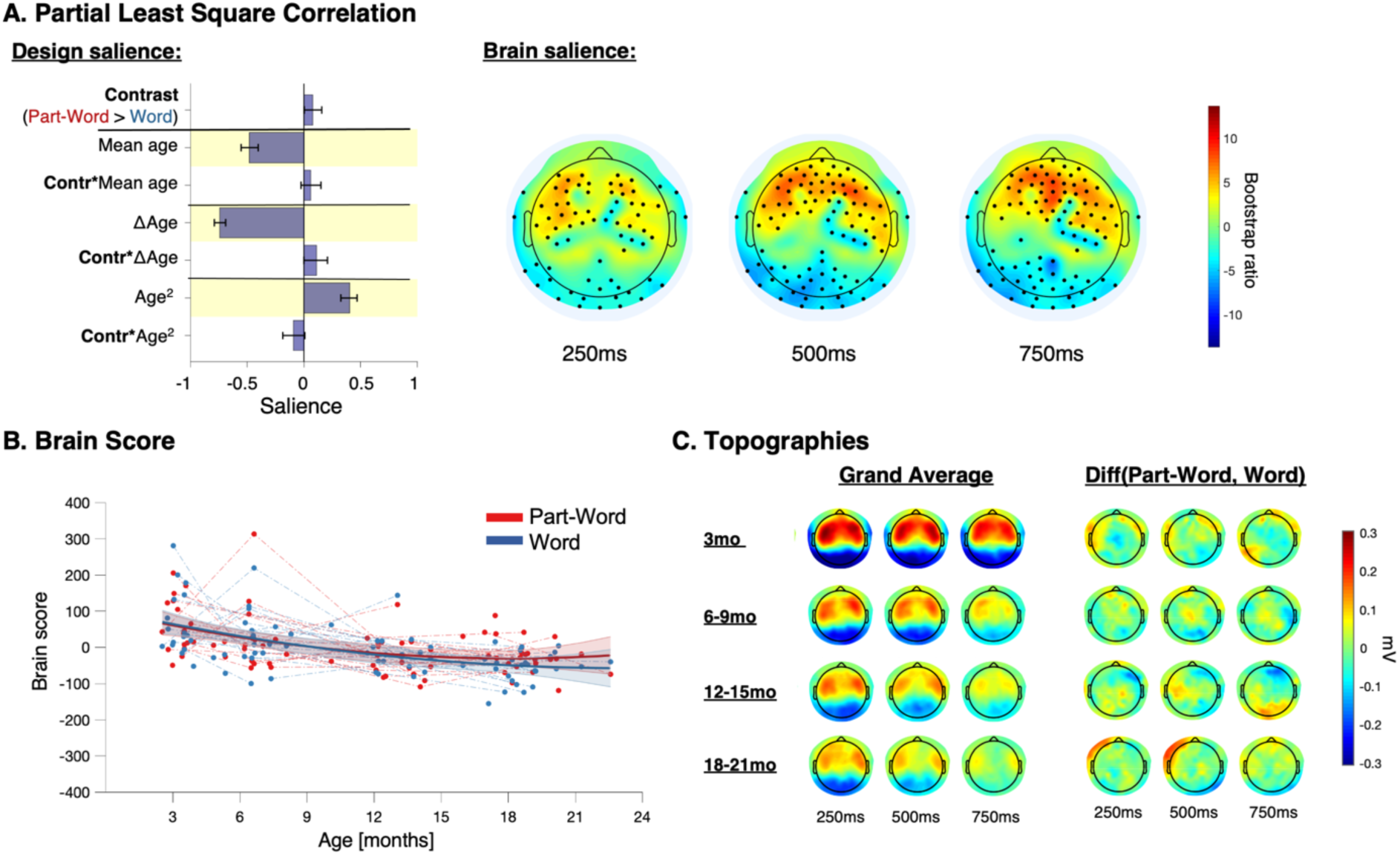
Early evoked response potential (ERP) to part-words compared to words across all participants. **A.** Design and brain saliences derived from the significant latent component. Brain topographies of bootstrap ratios (BSR) are displayed at 250ms intervals. Black dots indicate BSR > 2.3. **B.** Participants’ brain scores for part-word and word conditions, as a function of age. Brain scores are participants’ raw voltage data projected onto electrode saliencies. Brain scores illustrate how individual EEG data fit the saliences derived from the latent component. Linear fitting is used for illustrative purposes only. **C.** Voltage grand averaged (left) and differential (right) responses to part-word and word conditions at each age bin: 3 months (n=18), 6-9 months (n=20), 12-15 months (n=20), 18-21 months (n=25).

A PLS-c analysis of the differential ERP to part-words vs. words using group as contrast (HL *versus* LL) with age-related and verbal outcome parameters did not reveal any significant latent component. This indicates that the absence of an early condition effect in the full sample (**Figure 4**) was not due to opposing response patterns between the LL and HL groups.

### Late ERP

The PLS-C analysis in the late time window (1500 to 3000 ms), contrasting words vs. part-words, resulted in one significant latent component (p<.001, R=.49, 47.4% explained covariance) (**Figure 5**). The part-word vs. word contrast showed a significant contribution (BSR: 6.9), with a late frontal negativity and posterior positivity more pronounced in the part-word condition than in the word condition (see Supplementary materials, **Figure S6** for grand average topographies). All age variables contributed significantly to the latent component (BSRs for age: -10.9; delta-age: -21.6; age²: 8.6), with a similar pattern to that observed in the early time window: a decrease in grand average amplitude with age, following a slightly concave trajectory. Although the contrast × age² interaction reached statistical significance (BSR = –3.2), the corresponding effect (i.e., reduced condition differences at the youngest and oldest age) was modest and not clearly visible in the data (**Figure 5B-C**). Additional data will be needed to assess the robustness and replicability of this pattern. The same PLS-c in the sleeping subsample yielded no significant latent component, suggesting that sleep may reduce or even abolish the late response to word novelty.

**Figure 5.**
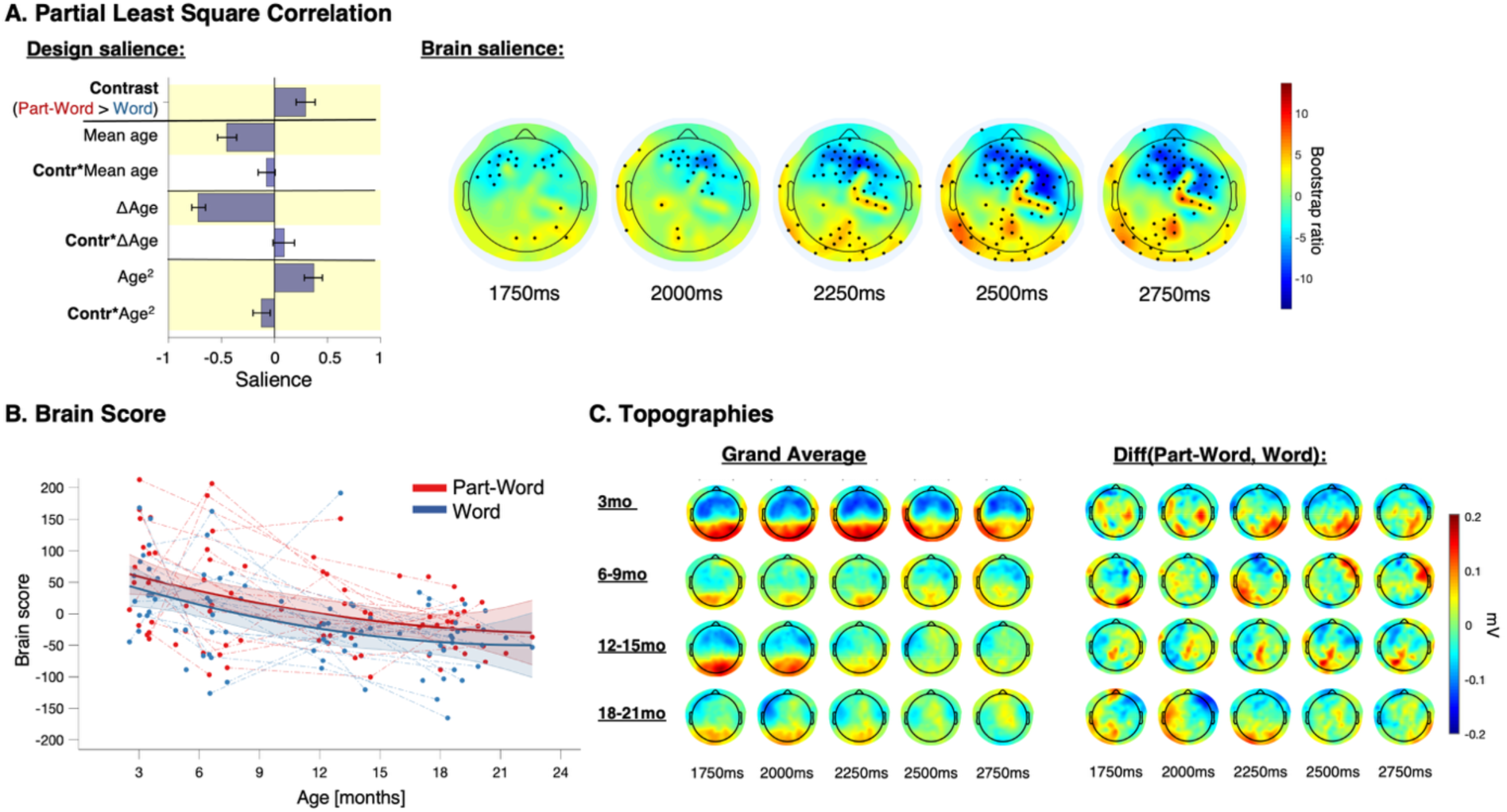
Late evoked response potential (ERP) to part-words compared to words across all participants. **A.** Design and brain saliences derived from the significant latent component. Brain topographies of bootstrap ratios (BSR) are displayed at 250ms intervals. Black dots indicate BSR > 2.3. **B.** Participants’ brain scores for part-word and word conditions, as a function of age. For details, see figure 4. **C.** Voltage averaged (left) and differential (right) responses to part-word and word conditions at each age bin: 3 months (n=18), 6-9 months (n=20), 12-15 months (n=20), 18-21 months (n=25).

A PLS-c analysis of the difference part-words *minus* words using group as contrast revealed one significant latent component (p=.002; r=.65; 29.3% explained covariance, **Figure 6**), with a similar topography and timing as the latent component identified on the activation using words vs part-word as contrast (Supplementary materials, **Figures S7-8** show grand average topographies per group and condition). This latent component included a significant group effect, with stronger frontal negativity and posterior positivity in LL participants compared to HL (BSR: 15.7). Additionally, significant negative age (BSR: -5.8), age² (BSR: -3.7), and age²*group interaction (BSR: -12.1) effects were observed. Furthermore, the latent component included a positive effect of both verbal outcome (BSR: 16.1) and verbal outcome*group interaction (BSR: 3.0). This indicates that the magnitude of the late response to novelty predicted verbal outcomes. Given that no late response was detected in sleeping participants, we re-ran the PLS-c analysis using group as a contrast in the awake subsample (n=58). This yielded one significant latent component (p=.006, r=.68, 30.5% explained covariance), with behavioral and electrode contributions highly overlapping with those in **Figure 6**.

**Figure 6:**
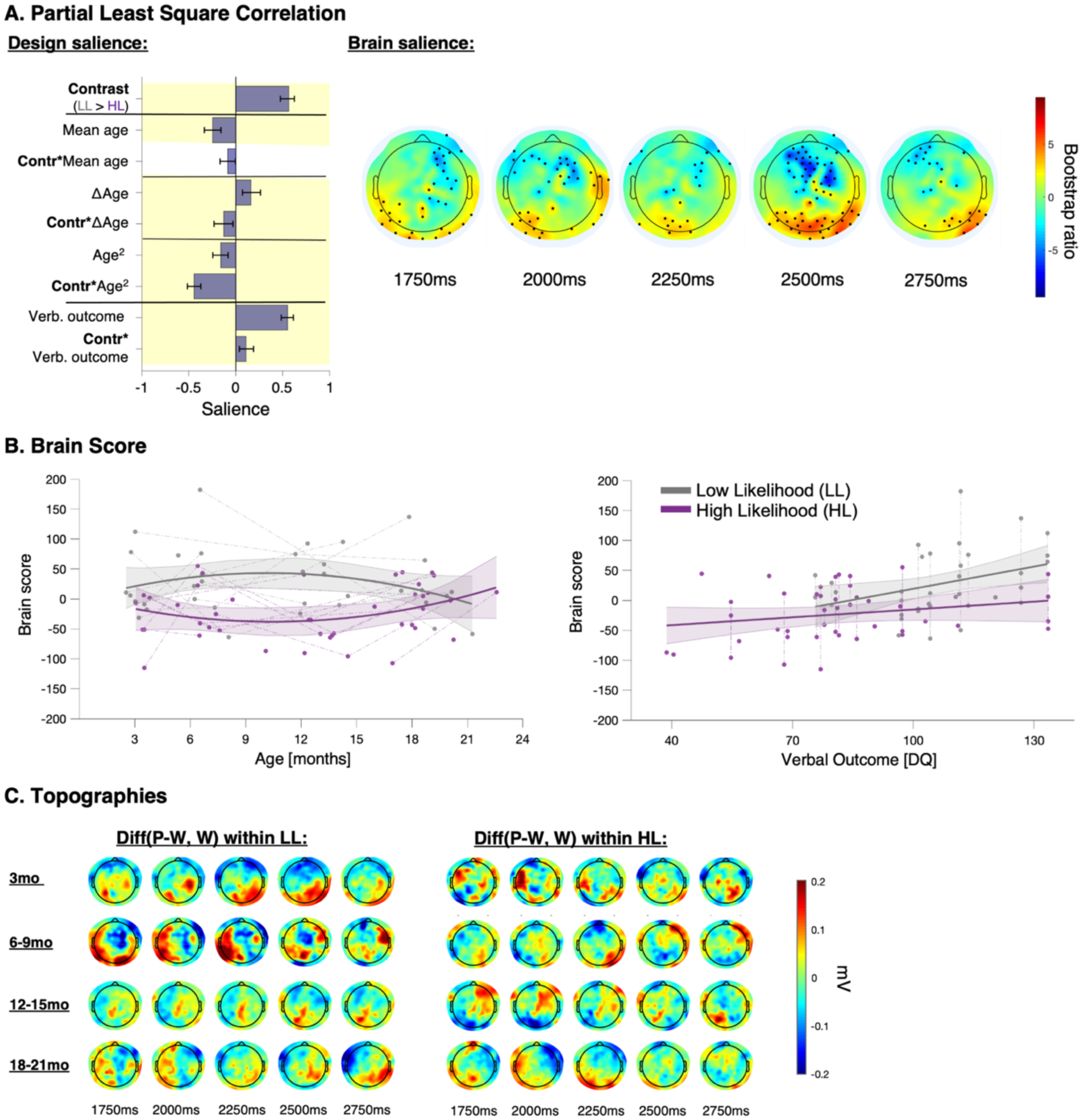
Late evoked response potential (ERP) to novelty in infants at high and low likelihood for autism. **A.** Design and brain saliences derived from the significant latent component. Brain topographies of bootstrap ratios (BSR) are displayed at 250ms intervals. Black dots indicate BSR > 2.3. **B.** Participants’ brain scores for part-word and word conditions, as a function of age (left) and of verbal outcome (right). Detailed information about brain scores is provided in Figure 4. Linear fitting is used for illustrative purposes only. **C.** Voltage differential responses to part-word and word conditions at each age bin and within each group: 3 months (11 LL and 7 HL), 6-9 months (9 LL and 11 HL), 12-15 months (10 LL and 10 HL), 18-21 months (9 LL and 16 HL). DQ: developmental quotient. HL: high likelihood for autism; LL: low likelihood for autism.

As we did for neural entrainment to syllables, we conducted a new analysis on late ERP to word novelty, excluding EEG data from the final visit. This PLS-c yielded one significant latent component (p=.002, r=.74, 33.5% explained covariance, **Figure 6 -figure supplement 1**) with globally similar EEG parameter contributions and age trajectory modelling. Verbal outcome still significantly contributed to the latent component (BSR: 20.1), with a negative verbal outcome*group interaction (BSR: -7.2). These results suggest that, after ruling out cross-sectional correlations at 18–21 months, the late ERP to word novelty predominantly predicts verbal outcomes in high-likelihood infants for autism.

Given that PLS-c analyses in the late window revealed both condition effects (words vs. part-words) and group differences on overlapping latent components, we asked whether the observed effect was present in both groups or driven primarily by LL participants. To test this, we ran separate supplementary PLS-c analyses within each group using the part-word vs. word contrast (see **Figure 6 -figure supplement 2-3**). Both analyses revealed a significant latent component (*p*<.001), with similar spatial topographies and temporal profiles. Crucially, the condition contrast contributed significantly only in the LL group (BSR: 13.0), but not in the HL group (BSR<2.3), suggesting that the group difference observed in **Figure 6** reflects the absence of a detectable novelty response in HL participants with the current analysis.

Overall, we found no difference between conditions during the early time window, which was aligned with the temporal progression of syllables within the triplet ([0–1000] ms; **Figure 4**). However, in a later time window ([1500–3000] ms) following triplet onset, the ERP exhibited a differential response to part-words and words (**Figure 5**). This late effect was positively associated with verbal outcomes at 20 months and, notably, was not detected in the HL group.

## Discussion

We exposed 44 infants at low (LL) and high (HL) likelihood for Autism Spectrum Disorder to an artificial speech stream composed of syllabic triplets (words) with random speaker changes at every syllable to assess their ability to track both syllables and word-level structure under conditions of acoustic variability. We focused on three electrophysiological measures: 1) neural entrainment to syllables; 2) neural entrainment to words; and 3) ERP responses to isolated words and part-words, analyzed in early [0-1000ms] and late [1500-3000ms] windows. These analyses allowed us to trace the sequential stages of speech processing, from syllabic perception to online word segmentation and later word recognition, and to identify the levels at which HL infants may encounter difficulties. While the core mechanism of statistical learning appeared intact in HL infants, as evidenced by word-level entrainment in both groups, several group differences emerged in the dynamics of syllable tracking and in the late ERP responses. Notably, some of these effects were significantly related to verbal outcomes at 18-21 months.

### Syllable neural entrainment

HL infants exhibited significantly weaker syllable-level neural entrainment than their LL peers (**Figure 2**), despite following a similar convex developmental trajectory across the first two years of life. This persistent lag in rhythmic tracking was not only stable over time but also strongly predictive of verbal outcome at 20 months, highlighting its potential as an early neurophysiological marker of language development. This result refines previous observations regarding reduced theta power in HL infants and autistic children exposed to natural speech (28,29). Unlike these studies, which estimated speech-brain coherence across the theta band, our analysis specifically targeted the particular frequency at which the auditory sequence was presented (4 Hz) and isolated phase-locked activity by subtracting adjacent frequency components. This more selective approach points to a precise impairment in synchronization with syllabic units, rather than a general reduction in low-frequency neural activity, indicating a targeted disruption of syllabic tracking.

Interestingly, the divergence between HL and LL infants emerged specifically during the structured stream and coincided with the time window in which infants began to segment the continuous speech flow into word-like units (Supplementary Material **Figure S4-5**). This temporal overlap suggests that successful segmentation may enhance syllable tracking via top-down predictions of the next syllable, improving alignment to syllable onsets in LL infants as well as in HL with better verbal outcome.

While this top-down interpretation highlights the role of word segmentation in enhancing syllable tracking, it does not exclude additional contributions from lower-level sensory mechanisms. According to the oscillatory framework of speech perception, syllable tracking is thought to be supported by endogenous neural oscillations in the theta band, which align with the natural rhythm of speech (54). Disruptions in this mechanism have been proposed in autism (28,29). However, recent evidence suggests that this model may not fully apply to early infancy. In typical 3-month-old infants, we observed robust neural entrainment to amplitude-modulated sounds across a wide frequency range (2–45 Hz), with more accurate tracking below 12 Hz — but no selective enhancement in the theta range (Kabdebon et al., submitted), challenging the idea of particular sensitivity to entrainment in this frequency range, at least in infants.

Beyond oscillatory alignment, another possibility is that weaker syllable tracking in HL infants stems from temporally and/or frequency-imprecise responses along the auditory pathway. Impaired temporal or spectral fidelity, particularly at the brainstem level, has been reported in autism and other language-related developmental disorders (37,55). In this context, decreased cortical power at specific frequencies observed in dyslexic adults (56) and autism (57) might reflect the long-term consequences of lower-level auditory processing difficulties rather than their root cause. Supporting this view, recent findings show that atypical brainstem responses can be detected even in first-degree relatives of autistic individuals, and that lower brainstem response consistency is associated with poorer pragmatic language skills in these individuals (58). These results highlight the cascading effect of low-level auditory processing anomalies on higher-level verbal and communication abilities. Such a cascade could explain the observed relationship between syllabic entrainment and verbal outcomes at 18–21 months. Future studies should directly assess whether degraded brainstem responses underlie reduced entrainment in HL infants using higher EEG sampling rates.

In sum, HL infants show a specific and developmentally stable deficit in syllable tracking, which may reflect reduced top-down support from word segmentation, impaired oscillatory alignment, and/or imprecise encoding in the brainstem. Crucially, this neural signature was predictive of later verbal outcomes, underscoring its potential relevance as both a mechanistic insight and an early clinical marker. We now turn to word entrainment results, which offer a complementary window into how infants begin to extract and consolidate word units from speech.

### Word segmentation: neural entrainment

Neural entrainment at the word rate (1.3 Hz) provides a robust index of speech segmentation: it can only emerge once infants begin grouping syllables into coherent word-like units. Despite the acoustic variability introduced by speaker changes, both HL and LL infants showed significant word-level entrainment after 90 seconds of exposure to the structured stream (Supplementary Material **Figure S3**), replicating previous findings with single speakers (15,16). Although HL infants exhibited reduced syllable tracking, no overall group difference was observed for word entrainment (**Figure 3**). However, the low frequency of the word rhythm, coupled with high power in the same range in infant EEG, may reduce the signal-to-noise ratio, limiting the sensitivity of this comparison.

Our longitudinal design, however, revealed a striking divergence in developmental trajectories. LL infants displayed a U-shaped curve in word entrainment, with strong responses at 3 and 21 months and a dip around 12 months (**Figure 3D**). In contrast, HL infants showed a flatter trajectory, with consistent performance across ages. Given the scarcity of longitudinal studies on statistical learning in infancy (42), these data provide novel insight into the developmental dynamics of word segmentation. Importantly, previous studies have shown that statistical learning is present from birth (14,15,44), and that word learning trajectories in 6-month-olds resemble those observed in adults (16), suggesting that this 12-month dip does not reflect a lack of statistical learning ability, but rather a transient disruption of an otherwise early and robust mechanism.

The dip in LL infants around 12 months coincided with an increase in syllable entrainment (**Figure 2B**), making it unlikely to result from reduced data quality or measurement artefacts. We propose, as a post-hoc hypothesis, that the random changes in speaker identity between syllables disrupted word segmentation. In typically developing infants, voice processing performance improves toward the end of the first year (59,60), alongside the accumulation of implicit knowledge about how speech is typically structured — notably, that words are usually produced by a single speaker and that they do not overlap prosodic boundaries. These emerging priors may constrain the segmentation process by restricting it to plausible word units, as has been shown for prosodic boundaries (61). In our paradigm, where speaker identity changed randomly between each syllable, this conflict may have temporarily disrupted this process, leading to a reduction in power at the word frequency.

Simultaneously, growing attention to voice-related features may have improved alignment to syllables, consistent with the convex concave shape of syllable entrainment. The 12-month dip in word tracking may also reflect the attentional cost of reorienting to each new voice (62,63), during a period when infants are intensely engaged in learning social cues. This interference appears to be transient: word segmentation recovers at later ages, potentially as infants become more efficient at managing speaker variability and less disrupted/interested by voice changes. This trajectory aligns with adult findings showing that word segmentation in continuous speech is context-sensitive, shaped not only by input statistics but also by the relative weight of different priors (12–50).

HL infants, on the other hand, did not show this transient disruption. In this group, word entrainment remained stable over time. To account for this unexpected finding, we followed up on the post-hoc hypothesis proposed above: a reduced sensitivity to social and vocal cues observed in HL infants may have spared segmentation abilities by limiting the interference introduced by speaker variability (64) . If this post-hoc hypothesis holds true, LL and HL infants would differ not in their intrinsic ability to learn statistical regularities *per se*, but rather in how they integrate or suppress competing cues (such as speaker changes) during the segmentation process. It is important to note, however, that the present study was not designed to isolate and evaluate the specific impact of speaker changes on word segmentation. Consequently, this interpretation remains speculative, and additional research is required to further address this question.

While HL infants may paradoxically benefit from a reduced sensitivity to conflicting social cues in our specific paradigm, this same attenuation could become a disadvantage in more naturalistic settings, where voice identity supports word recognition, such as attributing speech to the correct speaker during a multi-party conversation. Nonetheless, this interpretation remains speculative in the absence of a control condition with a consistent speaker. Future studies manipulating speaker variability will be essential to determine its causal role in shaping the divergent developmental trajectories observed between groups.

### Word recognition: ERP

In the test phase, we observed a robust late ERP response to novel pseudo-words compared to familiar triplets, emerging around 750 ms after word offset (**Figure 5**). This response reveals participants’ ability to recognize the words they have been exposed to during the stream. It belongs to the family of late slow-wave components observed in infants, which have been associated with higher-order novelty detection and orientation mechanisms (53). Crucially, the amplitude of this late response correlated positively with verbal outcomes at 20 months, suggesting that the ability to track and later recognize newly learned words may serve as an early predictor of language development. As with syllable entrainment, the late ERP to novel words primarily predicted verbal outcomes in high-likelihood (HL) infants.

Despite their transient drop in word tracking at 12 months, LL infants still exhibited a difference between words and part-words at test, indicating that they had successfully memorized the transitional probabilities between syllables. In other words, voice-related interference during the learning phase seems not to prevent participants from recognizing a familiar word form in test. Statistical learning can thus remain intact even when evidence for online segmentation is weak or absent — a dissociation previously reported in neonates and adults (43).

In contrast, HL infants showed no clear ERP difference between novel and familiar triplets (**Figure 6 -figure supplement 3**). Given that both groups showed similar word neural entrainment during learning, this absence is unlikely to reflect a failure in statistical learning itself. Instead, it may point either to a deficit in novelty orientation — a phenomenon previously described in autism and HL infants (65,66), or to a failure to explicitly recognize the novel part-word. A recent study in adults showed that implicit and explicit traces of statistical learning rely on separate consolidation mechanisms, with implicit learning being more robust (67). Interestingly, this dissociation between spared implicit versus impaired explicit statistical learning in autism has been previously proposed and discussed in the literature (68,69). According to these studies, autistic impairments in top-down attentional processes, such as social orienting — which are critical for bootstrapping language acquisition (70) — may result in a heightened dependence on bottom-up mechanisms, including implicit statistical learning.

Finally, we found no early ERP distinction between part-words and words in the [0–1000 ms] window following word onset (**Figure 4**). This contrasts with findings in neonates, who rely on the first-syllable to recognize recently learned words (15). This result suggests that by 3 months of age, infants may shift from relying on recognition of initial syllables to encoding entire word forms, a more mature form of lexical storage and retrieval. Alternatively, the topology of early responses might change during development, making the identification of these patterns difficult, given the relatively small number of participants tested at each age.

### Implications for early detection and intervention

Our results highlight several neurophysiological markers that may prove useful for early identification of infants at heightened likelihood for language difficulties or neurodevelopmental disorders. Importantly, these measures rely on passive EEG paradigms, making them accessible, non-invasive, and feasible even in very young or at-risk populations.

Syllable neural entrainment, which was consistently weaker in HL infants and predictive of later verbal outcomes, may serve as an early indicator of atypical speech tracking. This deficit may disrupt the ability to process and integrate phonetic and phonotactic cues critical for internalizing the rules of the native language. Moreover, interventions known to enhance auditory encoding — such as music-based training — have shown benefits for brainstem precision and language outcomes (71,72), and may prove particularly valuable in at-risk populations.

Likewise, the absence of a late ERP orientation response in HL participants may represent an early neural signature of altered attention to novelty that can be used both as a non-invasive predictor of language development and as a potential target for early intervention. This potential biomarker might nevertheless be modulated by participants’ sleep status, warranting careful consideration of vigilance state in future studies.

## Limitations

Several limitations of this study should be acknowledged. First, given the multidimensional nature of the EEG data and the longitudinal design, we relied on Partial Least Squares Correlation (PLS-c) analyses to identify latent components linking neural responses with experimental and developmental variables. While this multivariate approach was necessary to reduce data dimensionality and handle collinearity, it captures only shared variance across participants. As a result, PLS-c may miss more localized or subtle effects that do not align with the dominant latent structures. Thus, although our longitudinal design offers valuable insights into developmental trajectories, the uneven age distribution and limited number of repeated measures per infant constrain the interpretation of individual growth curves. More densely sampled longitudinal data would be needed to precisely model intra-individual changes. Second, although all participants were primarily exposed to French, we did not quantify additional language exposure, precluding any analysis of its potential moderator effects on statistical learning in our groups and age-trajectories. However, prior work has reported no effect of bilingualism on auditory triplet segmentation in children (73). Finally, while we interpret reduced syllable entrainment and diminished novelty responses in HL infants as potential early markers of later verbal outcomes, the predictive validity and autistic specificity of these neural indices should be confirmed in larger cohorts and across more diverse developmental profiles.

## Conclusion

Our study underscores the value of longitudinal neuroimaging—both online and offline—for unpacking the hierarchical processes underlying infant speech processing in both HL and LL populations. Even in typical development, these mechanisms remain poorly characterized. By mapping their developmental trajectories across the first two years of life, we revealed how early speech processing abilities are not static, but dynamically shaped by concurrent maturational changes.

Importantly, our results illustrate how subtle low-level differences—such as reduced syllable tracking or diminished novelty responses—can cascade into broader developmental outcomes. This supports a neuroconstructivist perspective (74), where early neural variability may help explain later divergence in language and communication, both in typical and atypical pathways.

## Materials and Methods

### Participants

Forty-four infants from the ongoing Geneva Autism Cohort were included during longitudinal visits, contributing to 83 EEG recordings (4,75). The open longitudinal design comprised four longitudinal visits: (1) at 3 months, (2) between 6 and 9 months, (3) between 12 and 15 months and (4) between 18 and 21 months. Verbal outcome was collected at the last visit (18-21 months). Longitudinal neuroimaging designs reduce within-subject variability by distributing the information across time (76,77). All participants were raised in primarily French-speaking environments, with French as the dominant language at home and daycare. The standardized developmental assessment Mullen Scale of Early Learnings was administered at the last visit to estimate participants’ verbal developmental quotient (DQ), our outcome measure (78). Participants’ developmental ages at the 18-21 months visit were divided by their chronological age to get developmental quotients (DQ) centered on 100. DQs avoid floor-effects by not truncating the very low-performing participants’ scores (79,80). The current study was approved by the University of Geneva’s Ethics Committee, and parents provided written informed consent. Sample characteristics are provided in **Table 1**.

**Table 1.** Sample characteristics. . Statistical comparison between LL and HL samples. For categorical variable, chi square (χ2) was applied. For continuous variables, we used two-tailed independent T tests. P values < .05 are highlighted in bold.

| MEASURE [mean (SD)] | Low-Likelihood | High-Likelihood | P-Val |
| --- | --- | --- | --- |
| Number of participants (n=44) | 19 | 25 |  |
| Number of EEG recordings (n=83) | 39 | 44 |  |
| 3 months EEG age (n=18) | 3.3±0.6 (n=11) | 3.5±0.3 (n=7) | .309 |
| 6-9 months EEG age (n=20) | 6.6±0.8 (n=9) | 7.1±1.3 (n=11) | .320 |
| 12-15 months EEG age (n=20) | 13.0±1.0 (n=10) | 13.5±1.4 (n=10) | .371 |
| 18-21 months EEG age (n=25) | 19.0±1.1 (n=9) | 18.7±1.6 (n=16) | .438 |
| Female biological sex | 4 (21.1%) | 12 (48.0%) | .066 ($\chi^2$) |
| Age at verbal outcome | 19.6±2.4 (n=16) | 19.7±2.0 (n=24) | .886 |
| Verbal outcome [DQ] | **104.3±16.2** | **77.9±22.2** | **.001** |

The 19 LL infants were healthy full-terms (>37 weeks of pregnancy) without any reported autism in their 1^st^ degree relatives. From the 25 HL infants, 18 had an older autistic sibling (n=18), which is known to be associated with a 18% prevalence of autism (81). The 7 other HL infants presented with early parental concerns for autism, based on parental report prior to enrollment. Their Autism Parent Screen for Infants (APSI) total score at their 18-21 months visit was 15.6±6.4, [8-22] range – a score greater than 8 reflecting a 63% positive predictive value for autism in HL populations (82). One of the HL participants with early parental concerns had a PACS1 mutation, a condition associated with a 37% autism prevalence (83). Moreover, 4 HL infants were born preterm (range: [31-36] gestational weeks), a condition associated with a 7% autism prevalence (84). Corrected age was used by subtracting the number of prematurity weeks from the chronological age (78,85).

There was no statistically significant difference (chi-square, p>.05) between HL and LL in either biological sex, visits’ repartition, or age (**Table 1**). Verbal outcome was missing for 4 participants (3 LL and 1 HL, 4 recordings) because of drop-out before the 18-21 months outcome visit. Those participants were excluded from analyses using verbal outcome as a parameter.

Of the initial sample (103 EEG recordings), 20 acquisitions from 16 participants (5 LL and 11 HL) were not included (19.4% data loss), due to too many crying and/or motion artifacts after visual inspection of the data (3 LL and 13 HL recordings), or to examiner’s acquisition error (2 LL and 2 HL recordings).

### Stimuli

Stimuli and procedure were based on Fló et al’s paradigm (15,44). Twelve syllables were synthesized with MBROLA (86) using French phonemes and phonotactic rules. Each syllable lasted 250 ms, had a flat intonation, and was produced by 3 male and 3 female MBROLA speakers. We built two different stream conditions: a structured stream (STR) and a random control stream (RND) by concatenating the syllables with speakers varying randomly from one syllable to the next. This created a rhythmic syllabic presentation at 4Hz in both streams. In the RND stream, syllables were presented randomly, maintaining a flat transition probability of 1/11 between syllables. In STR, syllables were organized into 4 tri-syllabic words presented in random order but without repetition, resulting in transition probabilities equal to 1 within words and a drop in transition probability to 1/3 at word boundaries. Words were paced at 1.33 Hz (1/(3 × 0.25s). We built two RND streams lasting 90 s each, one long STR stream lasting 180 s, and 6 short STR streams lasting 30 s each. Note that the voice dimension was totally orthogonal to the present design, each syllable being randomly produced by one of the six speakers. Thus, 1) the voice was not constant within a word and 2) there was acoustic variability between words (i.e. different voices produced the same syllables). This implies a voice normalization process to recognize the same syllables and words across different occurrences. More details about the stimuli are provided in the **Supplementary Material**, Stimuli section.

To explore infants’ word recognition after familiarization with the STR stream, we constructed isolated triplets corresponding to *words* and *part-words*. *Words* exactly matched the triplets used to build the STR streams (*A_i_B_i_C_i_* structure, letters being syllables from the *i*th learned word). *Part-words* included the 2 last syllables of a word *i*, with a random first syllable of another word *k* (*B_i_C_i_A_k_* structure). In a *part-word*, the transition probability between the two first syllables was 1, but these syllables were not in the correct position in the word. Furthermore, the transition probability between the second and the last syllable was 0.33 instead of 1. While a late ERP difference would signal that infants were sensitive to any of this information, an early difference would signal participants’ recall of the ordinal position of the syllables within the learned words (Flò et al, 2022).

### Data Collection

We collected high-density electroencephalograms (EEG) with a 128-electrode net (Electrical Geodesics, Inc) referenced to the vertex. The sampling frequency was 250 Hz. Participants were sitting on their parent’s lap, and a silent cartoon was presented to keep them quiet and still. Stimuli were played on a Bose® Companion 2 Series III at a 50cm distance with an intensity of 75dB. The cartoon was not time-locked to the auditory stimuli and varied across participants. The same examiner (MG) collected all data. When participants were restless and/or not interested in the cartoon, the examiner engaged them in quiet activities (e.g., staring at bubbles/toys). Some EEGs were recorded while participants were asleep (61.1% at 3 months, 40% at 6-9 months, 20% at 12-15 months and 4% at 18-21 months). There was no significant difference (p=.797) in sleeping status between HL and LL (linear mixed-effect model with repeated measures). Neural entrainment and statistical learning have been shown to be present in sleep in adults and neonates (15,16). The experimental procedure is detailed in **Figure 1A**.

### Data pre-processing

Data were first resampled to 300 Hz to get an integer sample number in each three-syllabic item, then band-pass filtered (0.2-40Hz). APICE preprocessing pipeline (https://github.com/neurokidslab/eeg_preprocessing) was applied using EEGLAB toolbox 2020.0 on Matlab® R2018b (87,88). In brief, bad segments of data were identified using algorithms detecting low correlation with other channels (usually due to non-functional channels) and outlier values for the signal amplitude or changes in the signal amplitude (typically due to motion artifacts), with a threshold of 2 (outliers are values bigger than two interquartile ranges away from the third quartile). A sample was considered to contain motion artifacts if more than 30% of the working electrodes were rejected. An electrode was considered bad if more than 30% of the free-of-artifacts samples were rejected. Short rejected periods (less than 100 ms) were corrected using target PCA. In segments containing less than 30% of the electrodes marked as bad artifact data were spatially interpolated using spherical spline interpolation. A careful data visual inspection was also carried out with manual removal of remaining bad electrodes and motion artifacts (done by MG). Independent Component Analysis (ICA) and the iMARA algorithm for component classification (89) were used to remove physiological noise. Finally, bad electrodes were interpolated using spherical splines.

#### A) Neural entrainment analyses

Given the construction of the streams with regular stimuli, we expected a neural entrainment at the stimuli specific frequencies (syllabic rate for the STR and RND streams and word rate when words were discovered in STR). This specific entrainment should not be observed during rest. We thus measured neural entrainment at 4 Hz (syllabic rate) and 1.3 Hz (word rate) in each period and each recording. We did so by considering non-overlapping epochs of 7.5 seconds respecting the chronological order of presentation (90). Epochs contaminated by artifacts were excluded from the analysis. On average, we kept 86% of data during RS (range of included epochs: [11-24]), 88% during RND ([12-24]), and 91% during STR ([32-48]). The number of included epochs did not differ between LL and HL groups, nor vary with age, suggesting an even data quality across groups and age (see Data quality section in Supplementary Material**, S1**).

Each epoch was average-referenced and normalized by dividing by the standard deviation computed across all electrodes and samples of each epoch. Denoising source separation (DSS) was applied to remove stimulus-unrelated activity using spatial filtering (91). Briefly, a PCA was first applied to the epoched data. Afterward, a DSS filter was applied on the 30 first components, and its 6 first components were kept. Eventually, a fast Fourier transform (FFT) was applied to the denoised epochs to estimate their phase-locking value (PLV). PLV estimates how much EEG activity is synchronized in phase with a specific frequency (1):

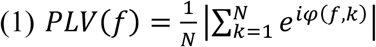

With N being the number of trials from one stream and 𝜑(𝑓, 𝑘) the phase at frequency *f* and trial *k*. PLV ranges from 0 (desynchronized activity) to 1 (phased locked activity). PLV was computed in 31 frequency bins (0.933 - 4.933 Hz, with 0.133 increment). In each stream, the PLV of each frequency bin was Z-scored over the 12 adjacent frequency bins (six on each side)(2):

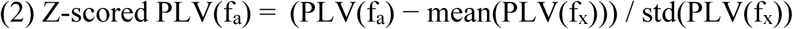

Where f_a_ is the targeted frequency bin, and PLV(f_x_) is the PLV over the 12 adjacent frequency bins. From now on, PLV refers to Z-scored PLVs. Additionally, in each electrode, PLV values from immediate adjacent frequencies (in which no entrainment is expected) were subtracted to get a cleaner signal (e.g., 1.2-1.47Hz for 1.3Hz word rate and 3.87-4.13Hz for 4Hz syllable rate).

### Longitudinal Partial Least Square Correlations

Partial Least Square Correlation (PLS-c) is a multivariate statistical approach that has been successfully implemented for EEG data (50,92,93). PLS-c can also be applied to longitudinal neuroimaging datasets, including a variable number of timepoints per participant, as ours (51,94). In a fully data-driven approach, one single longitudinal PLS-C can detect age-trajectories of electrophysiological measures (here, PLVs), indicate the electrode clusters in which those trajectories take place, and find their association with behavioral parameters (here, HL/LL group and verbal outcome). We applied longitudinal PLS-c based on the pipeline described in Delavari et al. 2021 using the myPLS toolbox (95,96) : https://github.com/danizoeller/myPLS). Briefly (**Figure 1B**): using each participant *i*’s *n^t^*^h^ visit, we built a behavior design matrix (Y^T^) with 9 variables: 1) Contrast (a binary contrast, in the figure example corresponding to LL or HL group), 2-3) two orthogonalized variables to grasp the cross-sectional and longitudinal effects of age (Mean age = averaged age across all *n* visits of the participant *i*, and Delta age = difference between participant *i*’s age at visit *n* and the participant *i*’s Mean age), 4) another age variable to grasp convex/concave trajectories (Age^2^=Delta age*[Mean age averaged across all participants]), 5-7) three interaction variables: Mean-age*Contrast; Delta-age*Contrast; Age^2^*Contrast, 8-9) Verbal outcome (DQ collected at the 18-21mo visit) and one interaction variable: Verbal outcome*Contrast. We limited the number of behavioral variables to nine to mitigate noise sensitivity and overfitting risks associated with exceeding the 10% sample size threshold (97). This limitation precluded the implementation of a single PLS-c model incorporating group, condition (STR vs. RND), age, and their interactions. Behavior design variables were z-scored across all participants. We also built a brain data matrix (X) for each participant *i*’s visit *n* including the 128 electrodes PLVs (1.3 Hz in STR when testing entrainment to word rate, and 4 Hz in both STR and RND when exploring entrainment to syllable rate). We then computed cross-variance matrices (R) as Y^T^X. R underwent singular value decomposition (R = USV^T^) to derive 9 singular values called latent components. Permutation testing (1000 permutations shuffling behavioral data across participants) estimated whether each latent component statistically significantly explained the correlation. Bonferroni correction was applied to account for multiple comparisons across the 9 tested latent components in the PLS-c, yielding an adjusted alpha of .006 (51,98). In each significant latent component, bootstrapping (500 random samples and replacement) was applied to evaluate the saliency of each behavior and brain (electrodes’ PLV) variable, reflecting the stability of its contribution to the latent component. Saliencies are summarized as bootstrap ratio (BSR): mean of bootstrap results divided by standard error. BSR are analogous to Z-scores and can be used to assess the stability of the saliency. We considered BSR > 2.3 as stable, corresponding to a 99.0% bootstrap confidence interval not crossing zero – roughly equivalent to a two-tailed p<.001 (50,51). To respect the intra-participant longitudinal dependencies, the bootstrap samples were randomly selected across participants and not EEG recordings. To help readers interpret the complex output of the PLS-c, we created plots of the individual raw EEG data extracted from the salient electrodes as a function of key behavioral variables (e.g., age, age-squared, and verbal outcome). We overlaid a polynomial curve (fitted using a least-squares mixed-effects model) on these raw data plots for visualization purposes only. This curve is strictly intended to assist with interpretation and is not meant as an additional statistical analysis, since the statistical relationship between EEG and behavioral variables was already established by the PLS-c.

It is important to note that PLS-c identifies EEG spatial patterns shared across ages and groups, allowing us to measure group and age effects within these shared spatial maps. Consequently, this method may overlook subtle spatial differences that could exist between specific groups and age categories.

#### B) Analyses of ERPs to test items

The preprocessed data were low-pass filtered (20 Hz) and epoched between [− 1.75, 3.25] s from the triplets’ onset. Epochs containing artifacts were excluded. On average, we kept 83% of data for Words (range of included epochs: [23-48]), and 83% for Part-Words ([19-48]). Included epochs tended to decrease with age, but did not differ between LL and HL groups, and there was no age*group interaction, suggesting a similar data quality across groups (Supplementary Material, **Figure S2**). Each participant’s data was reference-averaged and normalized by dividing by the standard deviation computed across all electrodes and samples of each epoch. Trials were averaged by condition (Words and Part-words).

### ERP longitudinal Partial Least Square Correlations

We applied longitudinal PLS-C on ERP data to investigate the trajectories of word recognition between LL and HL, using the same method as above with the following differences: brain matrices (X) included voltage measures from each electrode at each sample (3.33 ms time frame) instead of PLVs at each electrode. Consequently, X were time*space matrices (50). Moreover, to summarize the ERPs relationship with age and squared age, we computed brain scores. This metric reflects the projection of participants’ raw voltage values in the electrode saliencies obtained from the PLS. Brain scores thus provide summary values reflecting how well each EEG acquisition fits the brain saliences obtained by a given latent component. Fitting a mixed-effect model on brain scores allowed an illustration of the ERP trajectory with age. The same limitations outlined earlier apply to PLS-C when using ERP data — namely, it identifies spatial patterns shared between groups and age bins, which may lead to missing subtle, group- or age-specific differences.

## Contribution

A.F., G.D.L., M.G. and M.S. conceptualized the experimental protocol and analytic strategy. M.G. acquired and preprocessed the data. M.G. analyzed the data under the supervision of A.F. M.G. wrote the manuscript with the input of all authors. All authors read and approved the final manuscript.

## Acknowledgments

The authors would like to thank all the families who participated in the study, as well as the many collaborators who contributed to data collection over the years, especially Lylia Ben Hadid, Nada Kojovic, Kenza Latrèche, Sara Maglio, Irène Pittet, and Stefania Solazzo. We also would like to thank Farnaz Delavari for the invaluable pieces of advice provided for the statistical analyses.

---

**Figure 2 -figure supplement 1:**
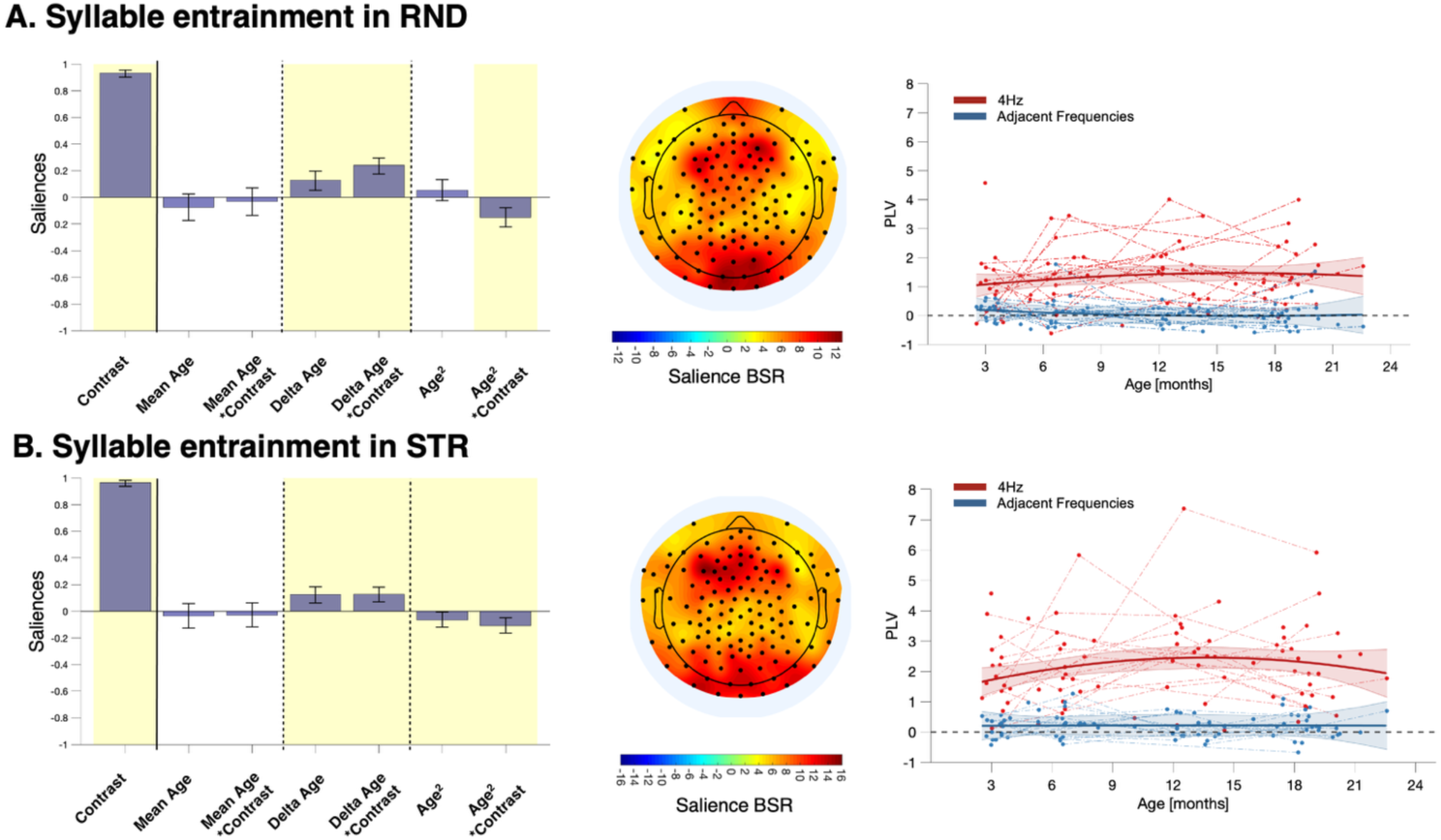
Follow-up analyses restricted to each stream for syllable entrainment. **A.** For RND, the latent component was significant (p<.001; r=.67; 87.3% explained covariance) with the following BSRs: contrast : 70.3*; mean age: -1.6; contrast*mean age: -0.7; delta age: 3.6*; contrast*delta age: 8.0*; age^2^: 1.2; contrast*age^2^: -4.1*. **B.** For STR, the latent component was significant (p<.001; r=.73; 91.1% explained covariance) with the following BSRs: contrast : 91.4*; mean age: -.9; contrast*mean age: 0.7; delta age: 4.2* contrast*delta age: 4.7*; age^2^: -2.4; contrast*age^2^: -3.8*. Yellow shading on left panels and black dots on middle panels indicate BSR > 2.3 – BSR >2.3 is considered significant.

**Figure 2 -figure supplement 2:**
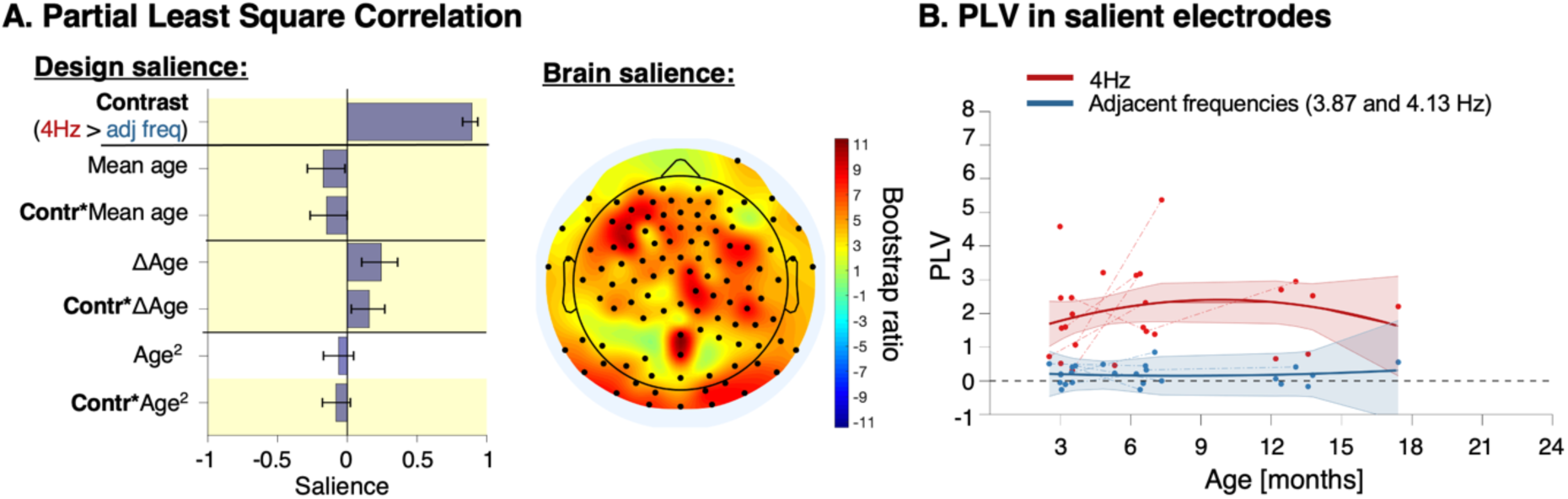
Neural entrainment to syllable rate in sleeping participants (n=25 recordings). **A.** Design salience (left) and brain salience (right topography) derived from the significant latent component for neural entrainment to the syllable rate using frequency (4hz versus adjacent frequencies) as contrast. Significance (i.e., salience) was established through bootstrapping. **B.** Individual raw Phase Locking Values (PLVs) extracted from the salient electrodes identified by the latent component (black dots in the topography on 2A) are displayed. A fitted curve is included for visualization purposes only, produced by a mixed-effects model with a 95% confidence interval.

**Figure 2 -figure supplement 3:**
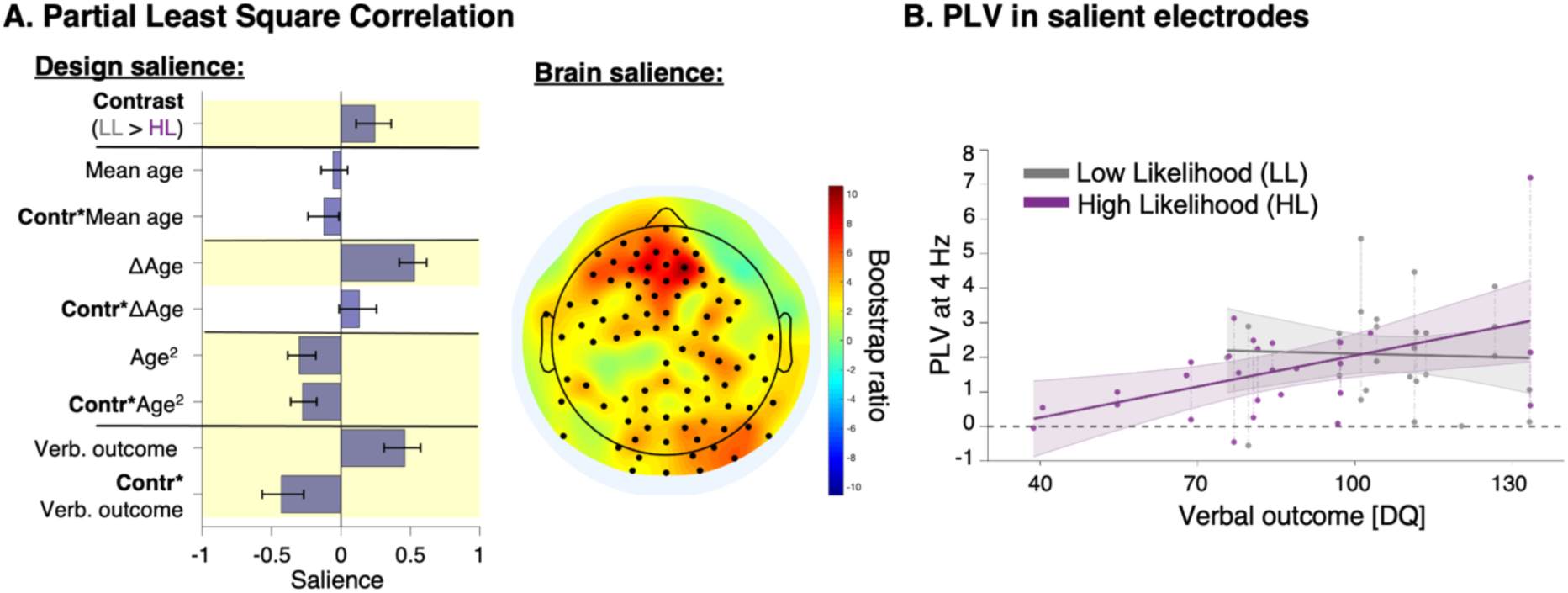
Neural entrainment to syllable rate (4 Hz), excluding the final visit (n=54 recordings). **A.** Design salience (left) and brain salience (right topography) derived from the significant latent component for neural entrainment to the syllable rate (4 Hz) using group (low versus high-likelihood for autism) as contrast. Significance (i.e., salience) was established through bootstrapping. Bars represent the mean of 500 random salience samples with replacement bootstrapping, and error bars indicate the 95% confidence interval. Yellow shading highlights variables that significantly contribute to the latent component, defined by a Bootstrap Ratio (BSR; mean of bootstrapping divided by standard deviation) > 2.3. The topography of the BSR values shows electrodes significantly contributing to the component (indicated by black dots, BSR > 2.3). **B.** Individual raw Phase Locking Values (PLVs) extracted from the salient electrodes identified by the latent component (black dots in the topography on 2A) are displayed. At every age, a verbal developmental quotient (DQ) of 100 is expected in the general population. A fitted curve is included for visualization purposes only, produced by a mixed-effects model with a 95% confidence interval. This curve is intended solely to aid visualization, as the statistical relationships between EEG and behavioral variables are determined by the PLS-C analysis.

**Figure 3 -figure supplement 1:**
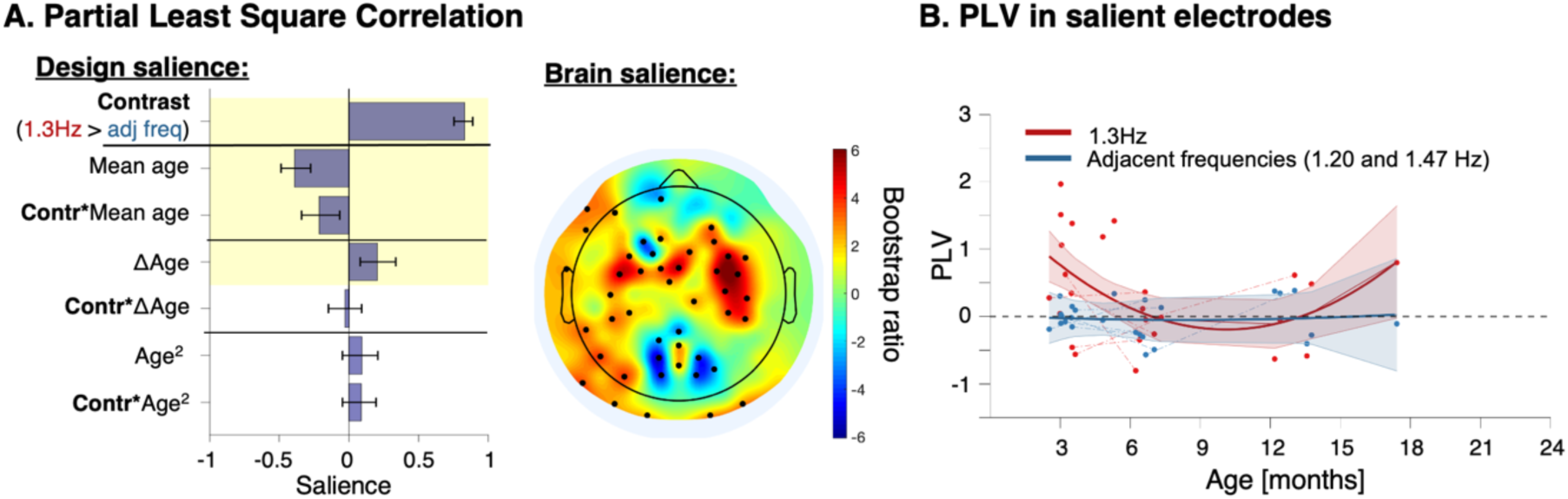
Neural entrainment to word rate in sleeping participants (n=25 recordings). **A.** Design salience (left) and brain salience (right topography) derived from the significant latent component for neural entrainment to the word rate using frequency (1.3hz versus adjacent frequencies) as contrast. Significance (i.e., salience) was established through bootstrapping. **B.** Individual raw Phase Locking Values (PLVs) extracted from the salient electrodes identified by the latent component (black dots in the topography on A.) are displayed. A fitted curve is included for visualization purposes only, produced by a mixed-effects model with a 95% confidence interval.

**Figure 3 -figure supplement 2:**
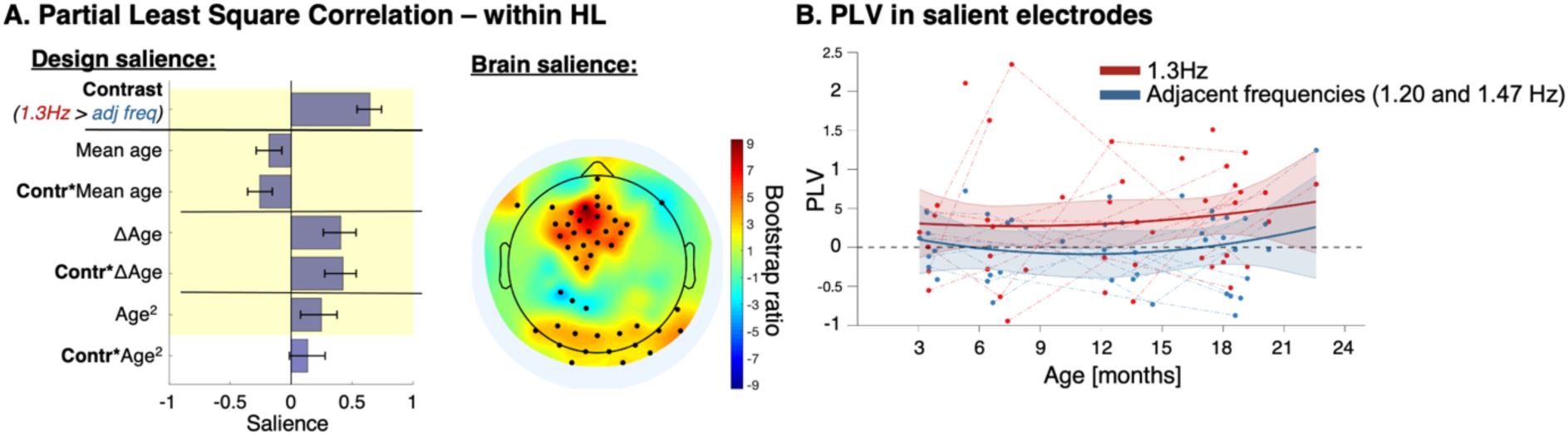
Word entrainment within HL participants (n=25; 44 recordings). **A.** Design salience (left) and brain salience (right topography) derived from the significant latent component for neural entrainment to the word rate using frequency (1.3hz versus adjacent frequencies) as contrast. Significance (i.e., salience) was established through bootstrapping. BSRs: contrast : 12.9*; mean age: -3.2*; contrast*mean age: -5.3*; delta age: 5.9*; contrast*delta age: 6.7*; age^2^: 3.6*; contrast*age^2^: 1.9. **B.** Individual raw Phase Locking Values (PLVs) extracted from the salient electrodes identified by the latent component (black dots in the topography on A.) are displayed. A fitted curve is included for visualization purposes only, produced by a mixed-effects model with a 95% confidence interval.

**Figure 6 -figure supplement 1:**
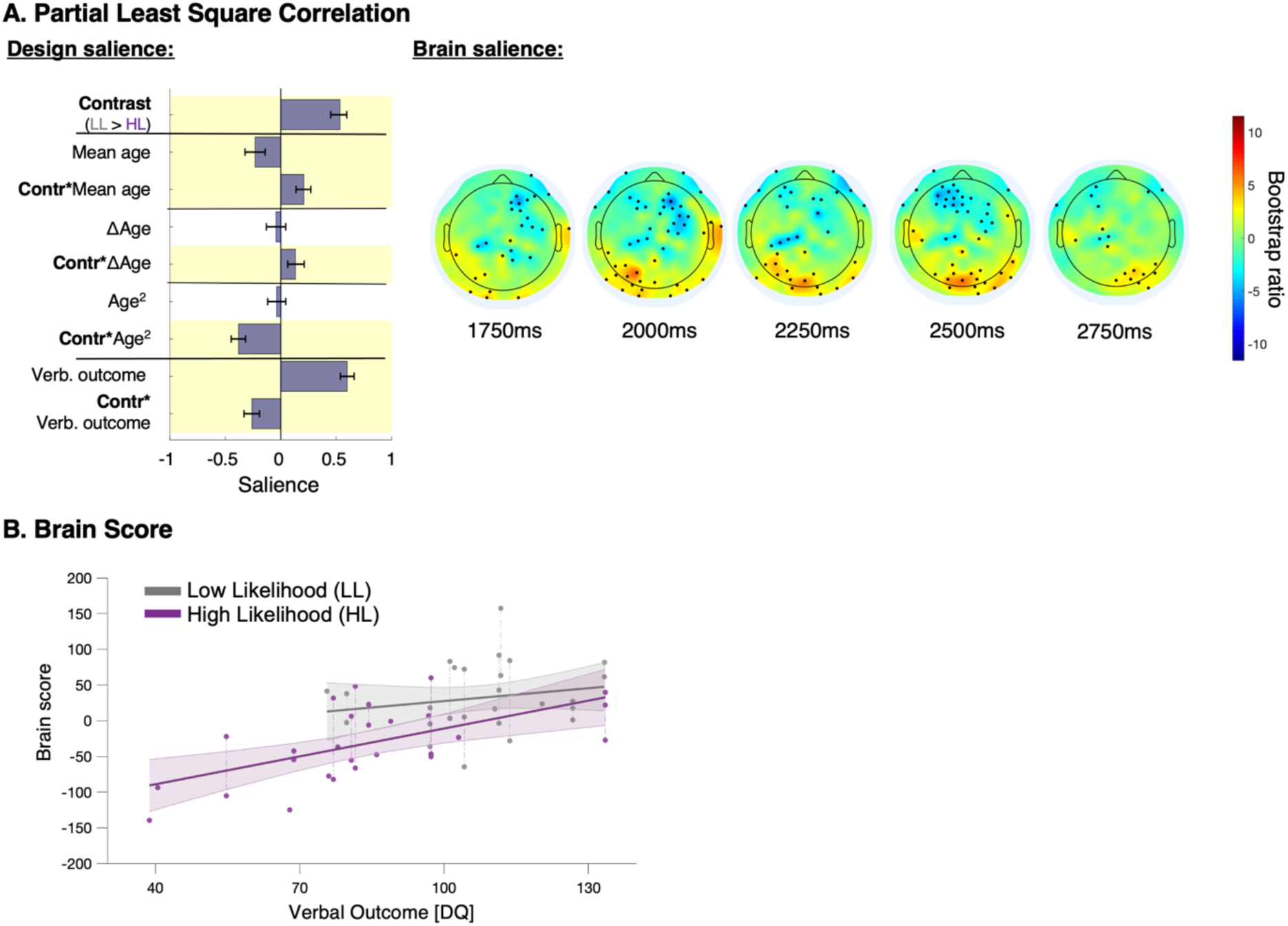
Late evoked response potential (ERP) to word novelty, excluding the final visit (n=54 recordings). **A.** Design and brain saliences derived from the significant latent component. Brain topographies of bootstrap ratios (BSR) are displayed at 250ms intervals. Black dots indicate BSR > 2.3. **B.** Participants’ brain scores for part-word and word conditions, as a function of verbal outcome. Brain scores are participants’ raw voltage data projected onto electrode saliencies. Brain scores illustrate how individual EEG data fit the saliences derived from the latent component. Linear fitting is used for illustrative purposes only. HL: high likelihood for autism; LL: low likelihood for autism.

**Figure 6 -figure supplement 2:**
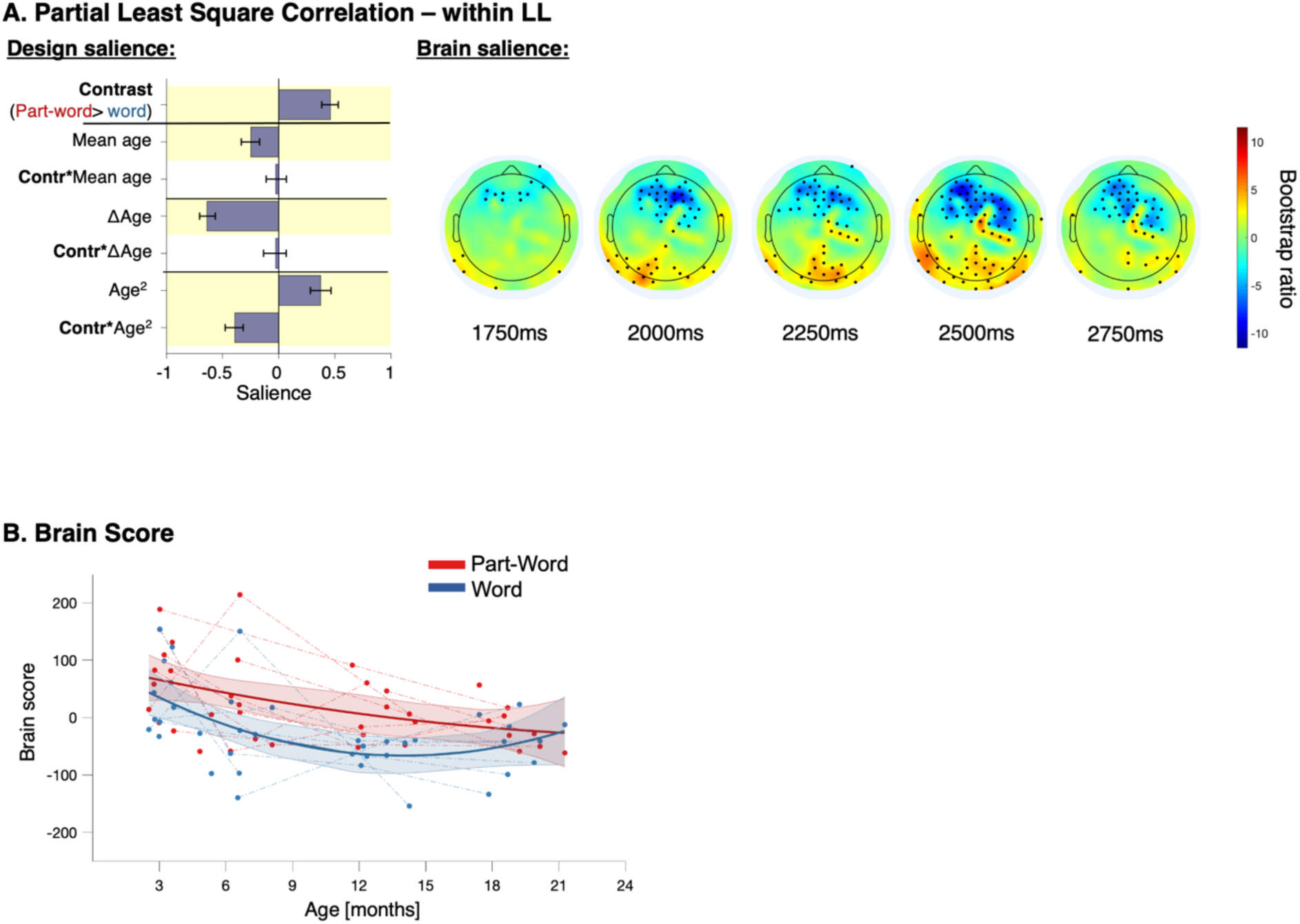
Late evoked response potential (ERP) to word novelty within low likelihood infants (n=19; 39 recordings) **A.** Design and brain saliences derived from the significant latent component. Brain topographies of bootstrap ratios (BSR) are displayed at 250ms intervals. Black dots indicate BSR > 2.3. **B.** Participants’ brain scores for part-word and word conditions, as a function of age. Brain scores are participants’ raw voltage data projected onto electrode saliencies. Brain scores illustrate how individual EEG data fit the saliences derived from the latent component. Linear fitting is used for illustrative purposes only. LL: low likelihood for autism.

**Figure 6 -figure supplement 3:**
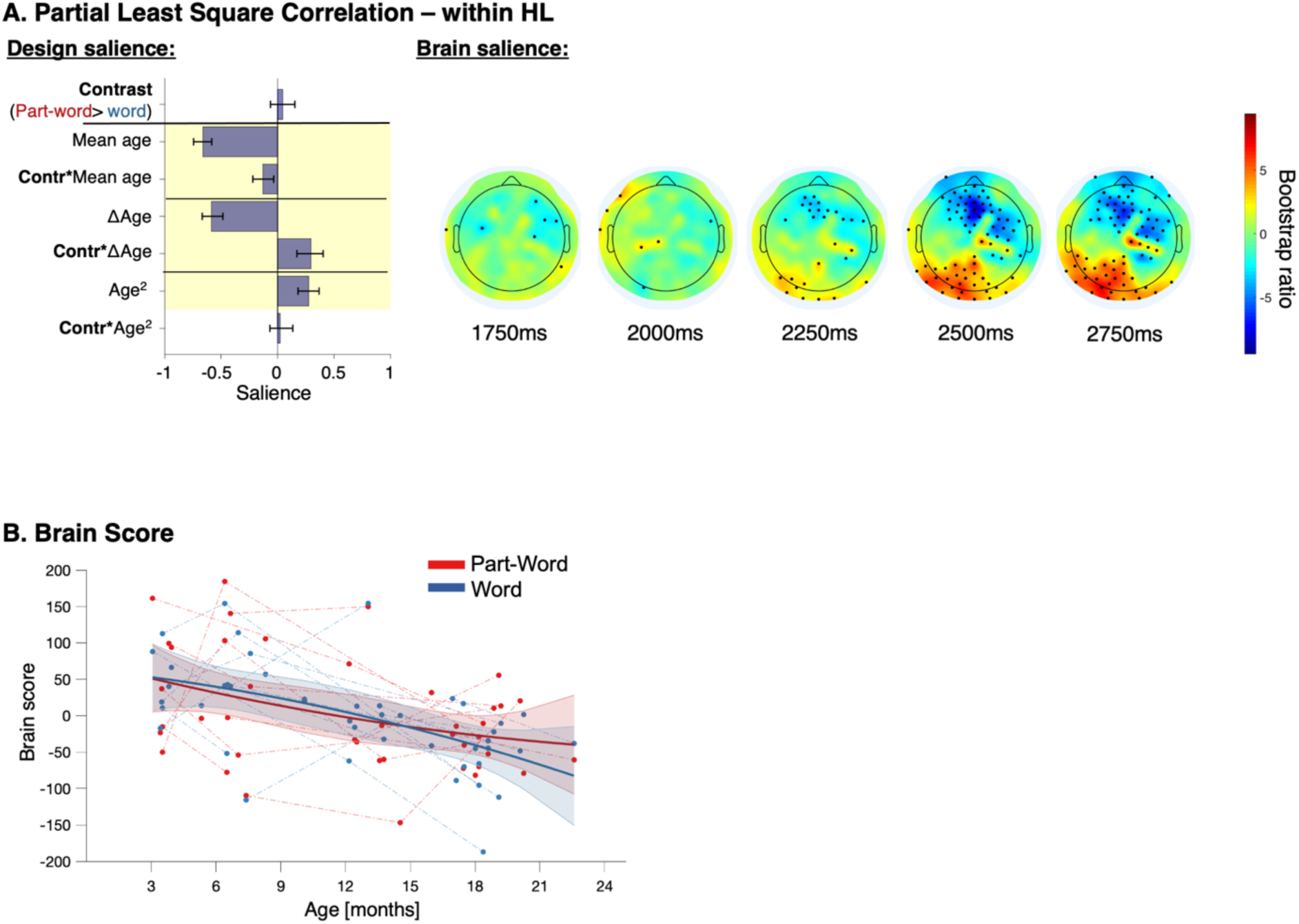
Late evoked response potential (ERP) to word novelty within high likelihood infants (n=25; 44 recordings) **A.** Design and brain saliences derived from the significant latent component. Brain topographies of bootstrap ratios (BSR) are displayed at 250ms intervals. Black dots indicate BSR > 2.3. **B.** Participants’ brain scores for part-word and word conditions, as a function of age. Brain scores are participants’ raw voltage data projected onto electrode saliencies. Brain scores illustrate how individual EEG data fit the saliences derived from the latent component. Linear fitting is used for illustrative purposes only. HL: high likelihood for autism.

## Supplementary Material

### 1) Stimuli

Using the MBROLA diphone, we synthetized 12 syllables (250 ms duration) using French phonemes: 6 vowels with 160ms duration ([i], [e], [ɛ], [a], [u], [o]) and 12 consonants with 90ms duration, 6 voiced ([b], [d], [g], [v], [z], [ʒ]), 6 unvoiced ([p], [t], [k], [f], [s], [ʃ]). Syllables were [ta], [do], [vɛ], [fi], [za], [pu], [ge,], [kɛ], [so], [ʒu], [ʃe], and [bi]. All syllable combinations followed French phonotactic rules. Each syllable was produced by six selected speakers in MBROLA: 3 male adult speakers (fr3 with low pitch, fr1 with middle pitch, fr7 with high pitch) and 3 female adult speakers (fr2 with low pitch, it4 with middle pitch and fr4 with high pitch). Random (RND) and structured (STR) streams were constructed by concatenating the syllables without coarticulation and a random choice of the speaker, with the only constraints that there was no repetition of the same speaker in a row, nor alternation between two speakers more than once (if A and B are two speakers, then ABAB is forbidden). The same rule was applied for syllable identity in the RND stream and the word in the STR stream (i.e. no repetition, nor alternation more than once of a syllable or a word). Three STRs were built, each one using different words, and randomized across participants. The words in stream 1 were [tadovɛ], [fizapu], [gekɛso], and [ʒuʃebi], and part-words (including the 2 last syllables of a word with a random first syllable of another word) were [dovɛfi], [ʃebita], [bitado], and [soʒuʃe]. In the second stream words were [dovɛfi], [zapuge], [kɛsoʒu], and [ʃebita]. Part-words were [vɛfikɛ], [soʒuʃe], [tadovɛ], and [gekɛso]. In the third stream, words were [vɛfikɛ], [soʒuʃe], [pugeza], and [bitado]. Part-words were [tadovɛ], [ʒuʃebi], [kɛsoʒu], and [zapuge].

### 2) Data quality

#### A) Data quality for neural entrainment measures

We applied a linear mixed-effect model (MyMixedModelsTrajectories toolbox in Matlab R2018: https://github.com/danizoeller/myMixedModelsTrajectories) on individuals’ amount of included epochs in RS, RND and STR (total of included epochs, the maximum possible value being 96), with fixed effects for age and group (HL versus LL), and random slope that varied by participant. The number of included epochs for each participant was used as a proxy for data quality. Three random-slope mixed-effect models with random slopes (constant, linear, quadratic) were fitted using the following formula:

(1) Good-epochs ∼ age * group + (1+age|subject)

The following equation was used for the linear model:

(2) Good-epochs_im_ = β0 + β1*group_i_ + β2*age_im_ + β3*group_i_ *age_im_ + b1_m_* age_i_ + b0_m_+ ε_im_

for participant *i* at timepoint *m*, with β1-3 being the fixed-effect coefficients for group (coded by a dummy variable), age and group*age interaction. The b1_m_ term is the random slope varying by participant, b0_m_ is the normally distributed random effect, and ε_im_ is the normally distributed observation error. The Bayesian information criterion (BIC) indicated the best-fitting model. Here, the constant model was selected (BIC=584), suggesting that age had no effect on data quality. We found no significant group effect (p=.748, beta=85.5). Consequently, we considered that data quality for neural entrainment measures was broadly similar between HL and LL and across ages. This analysis is illustrated in the **Figure S1**.

**Figure S1:**
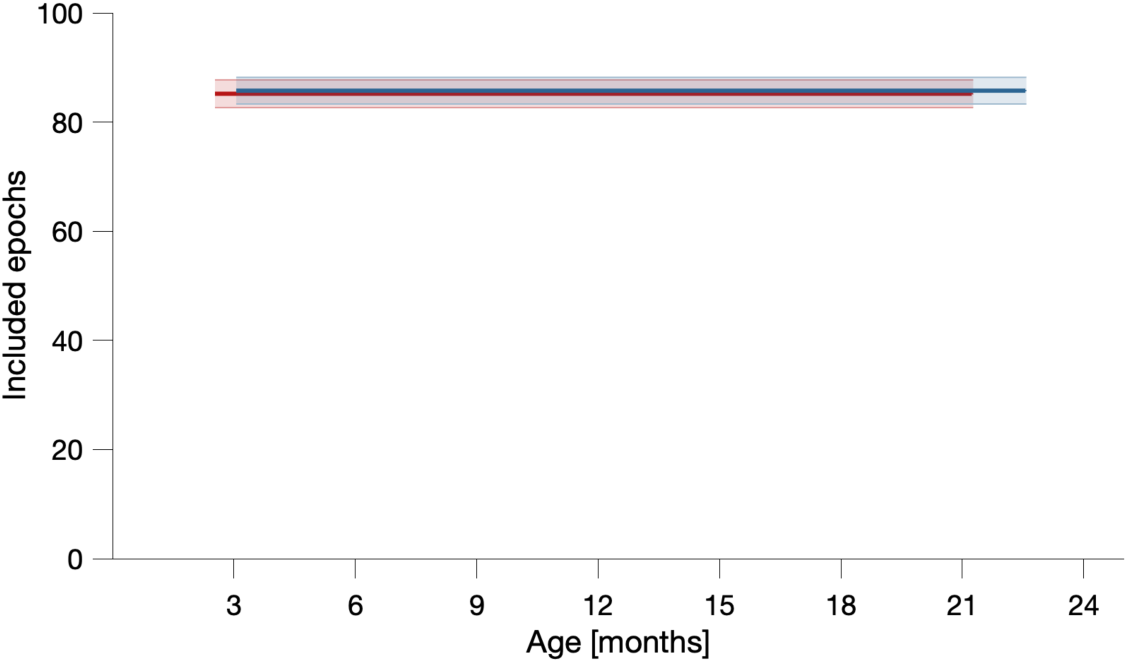
Neural entrainment data quality. LL mean trajectory is in red and HL in blue, with their respective 95% confidence intervals. Included epochs in absolute number.

#### B) ERP data quality

The same analysis was run on the ERP epochs (total of included epochs, the largest possible value being 96: 48 Word items and 48 Part-Word items). The model of order 1 (linear) was selected (BIC=569), we found no significant group effect (p=.656, beta = -1.6) or group*age interaction effect (p=.436, betas = -0.4) or age effect (p=.149, beta = -0.6). This analysis is illustrated on the **Figure S2.**

**Figure S2:**
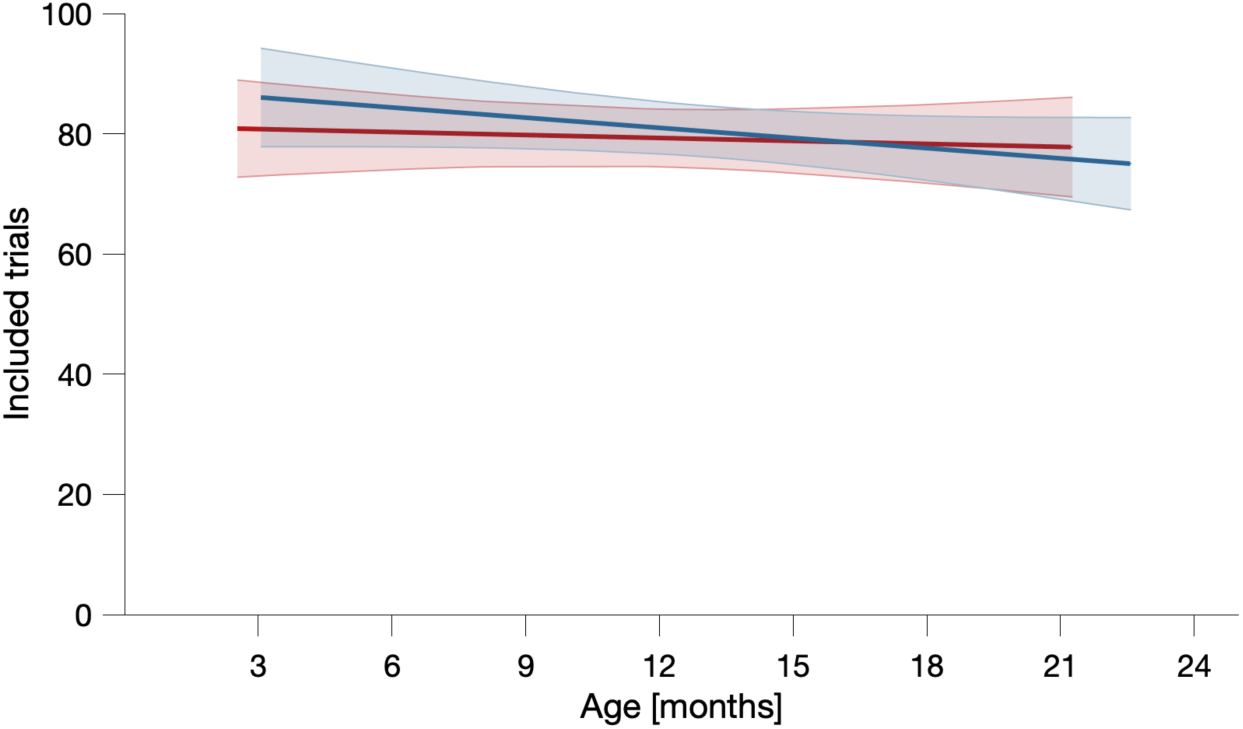
ERP data quality. LL mean trajectory is in red and HL in blue, with their respective 95% confidence intervals.

### 3) Supplementary analyses for neural entrainment

#### Learning time course across all participants

Additionally, we ran an analysis of neural entrainment over the time course of the stimuli presentation to provide an illustration of neural entrainment over the experimental session, as done in Smalle et al (2022) and Flò et al (2022) for instance. We concatenated the epochs in chronological order (180 seconds of random stream, 360 of structured stream, and 180s of random stream again). For each targeted frequency (word at 1.33 Hz and syllable at 4 Hz), we averaged PLVs over the salient electrodes obtained from the PLS-C (black dots from **Figure 3C** for 1.3Hz entrainment and from **Figure 2C** for 4Hz). For both frequencies, we fitted a mixed-effect model using Matlab *fitlme* function (PLV∼1+(1|participant)), using 120 second sliding windows with an increment of 1.5 second and testing for differences with zero (**Figure S3)**.

**Figure S3:**
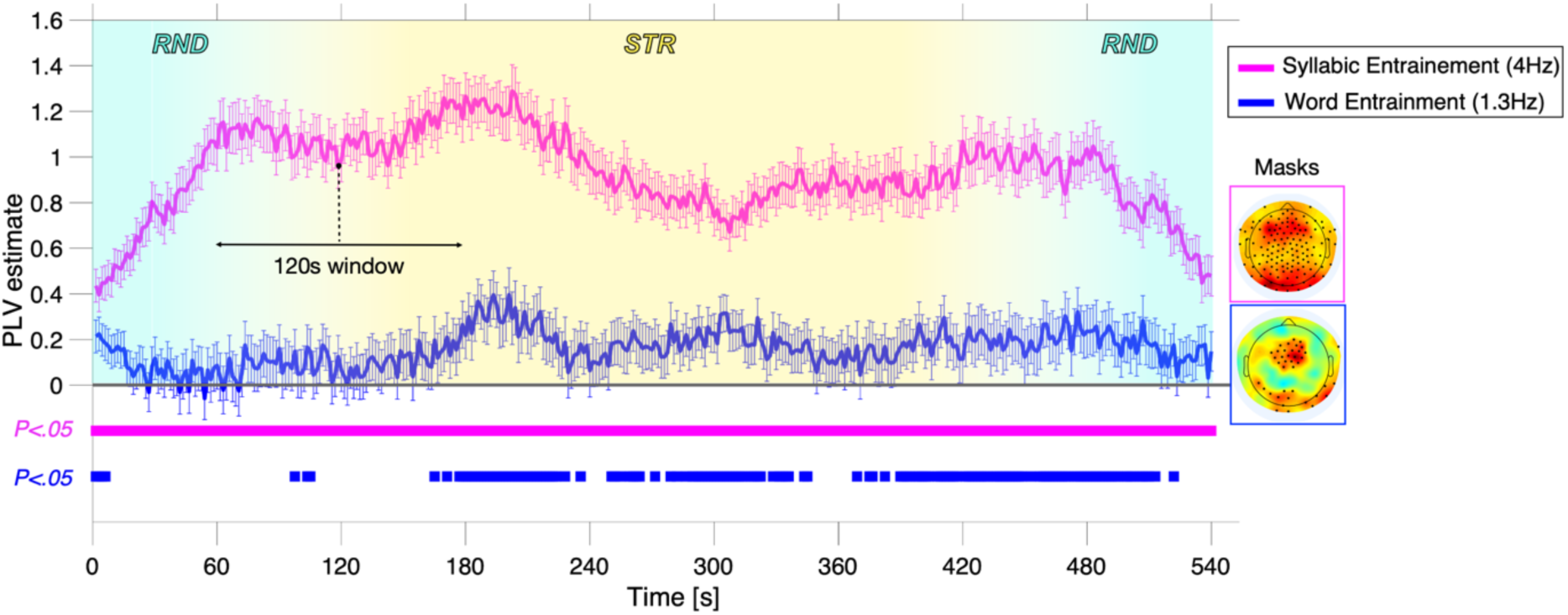
Neural entrainment time course over the experimental session considering all participants. The plain squares under the plots correspond to the time samples with PLVs significantly greater than 0 (p<.05).

#### Group differences in the time course of neural entrainment

We ran two PLS-cs to analyze this time dimension. Brain matrices included PLVs at each timeframe (120s time-windows with 1.5s increment) averaged in space across salient electrodes obtained from the PLS in space dimension (black dots on **Figure 3C** for word entrainment and black dots of **Figure 2C** for syllable entrainment). Design matrices were the same as in previous analyses (including group LL>HL as contrast and age-related variables and verbal outcome as factors).

The PLS-c for 4Hz syllable entrainment resulted in one significant latent component (p=.001; r=.66; 32.0% explained covariance) with the following BSRs: contrast (LL > HL): 7.2*; mean age: -12.3*; contrast*mean age: -1.4; delta age: 5.9*; contrast*delta age: -2.7*; age2: -1.8; contrast*age2: 2.2; verbal outcome: 20.5*; contrast*verbal outcome: -12.8* (**Figure S4**). PLS-c for 1.3Hz word entrainment gave no significant latent component (most significant latent component : p=.103) revealing no difference between HL and LL participants.

**Figure S4:**
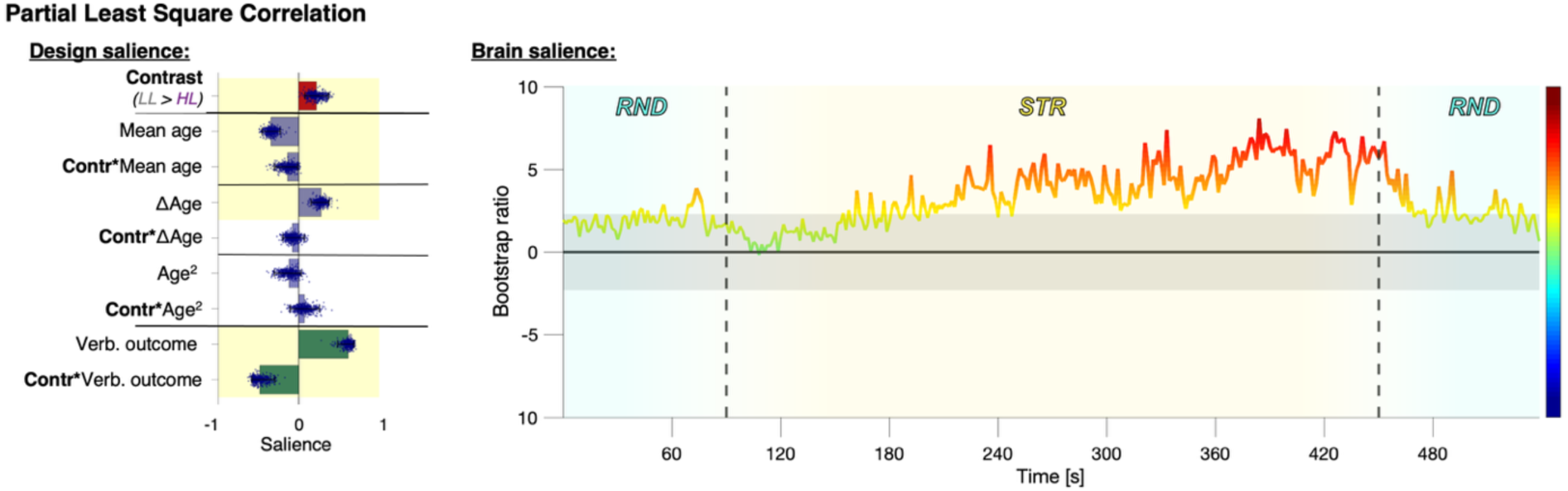
Group differences in the time course of syllable neural entrainment (4Hz). Gray shading on right panel indicates BSR < 2.3; BSR >2.3 is considered significant. Vertical dashed lines indicate the transitions between random and structured streams.

To better disentangle how each group tracks syllables in STR in contrast to RND, we ran a PLS-c testing group effect on 4hz PLVs in each stream. In the RND stream, the analysis yields one significant component (p<.001, r=.49, 52.7% explained covariance, **Figure S5A-B**). Bootstrap ratios (BSR) were: contrast (low versus high likelihood) 6.9; mean age -1.0; contrast*mean-age 0.1; delta-age 11.8; contrast*delta-age 0.1; age^2^ -4.9; contrast*age^2^ 4.6; verbal-outcome 9.7; contrast*verbal-outcome -0.6. The model still highlighted a strong link between syllable tracking and group, suggesting that RND also discriminates between HL and LL. However, RND syllable tracking doesn’t appear to be linked to group x verbal-outcome as we observed in **Figure 2C-D**. ). In the STR stream, we obtained one significant latent component (p=.002, r=.51, 57.8% explained covariance, **Figure S5C-D**). Bootstrap ratios (BSR) are: contrast 2.2; mean age -1.6; contrast*mean-age -1.1; delta-age 6.1; contrast*delta- age -0.7; age^2^ -4.4; contrast*age^2^ 0.5; verbal-outcome 8.2; contrast*verbal-outcome -7.3. Here, the strong association between syllable tracking and group*verbal-outcome is similar to the model presented in **Figure 2C-D**.

**Figure S5:**
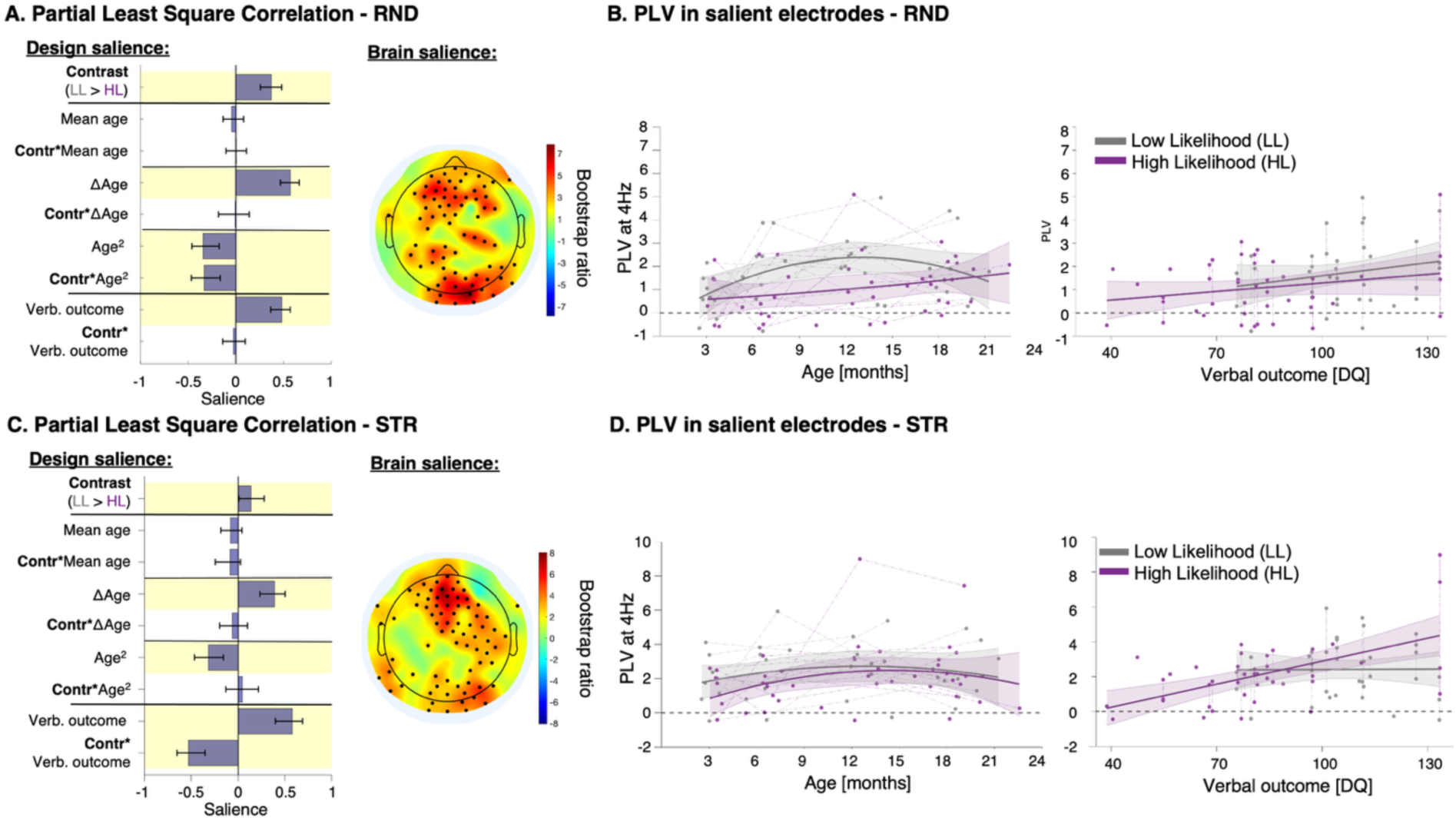
Syllable entrainment within RND (A-B) and STR (C-D).

### 4) Supplementary analyses for ERP

First, ERP topographies were inspected across ages in the whole sample to exclude the presence of maturational qualitative changes that would have prevented the inclusion of all recordings into the same longitudinal PLSC model. **Figure S6** shows the ERP topographies averaged across all participants at each visit for each condition (Part-word and Word), as well as the difference between conditions (Part-Word minus Word). No salient maturational qualitative change emerges from visual inspection (e.g., polarity inversion), legitimizing the use of a single multivariate linear model (PLS-C) that includes all visits.

**Figure S6:**
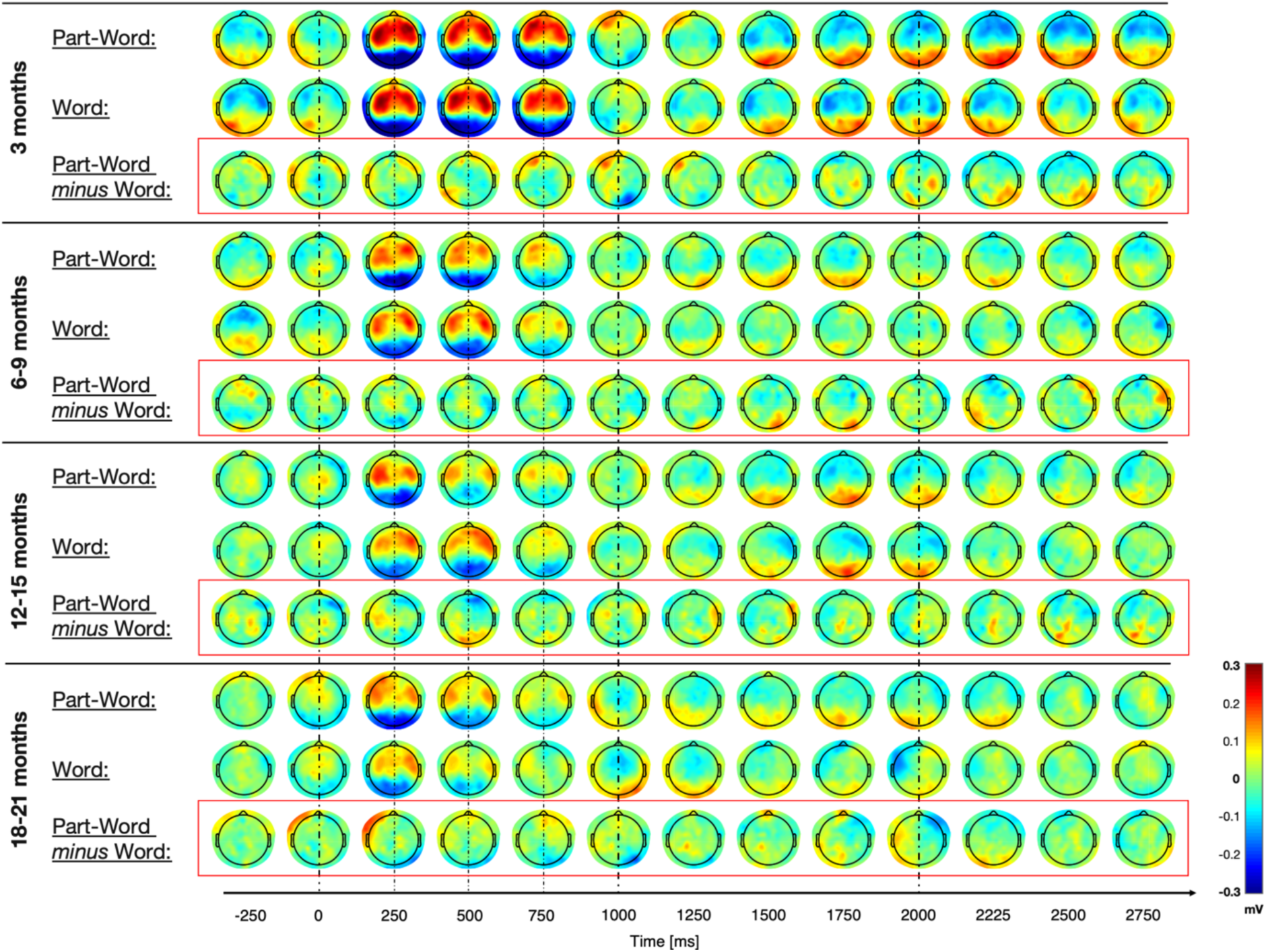
ERP topographies across age bins and participants.

**Figures S7-8** further divide ERP topographies by group, showing the response in both LL and HL at each age bin and for each condition: 3 months (n=18), 6-9 months (n=20), 12-15 months (n=20), 18-21 months (n=25).

**Figure S7:**
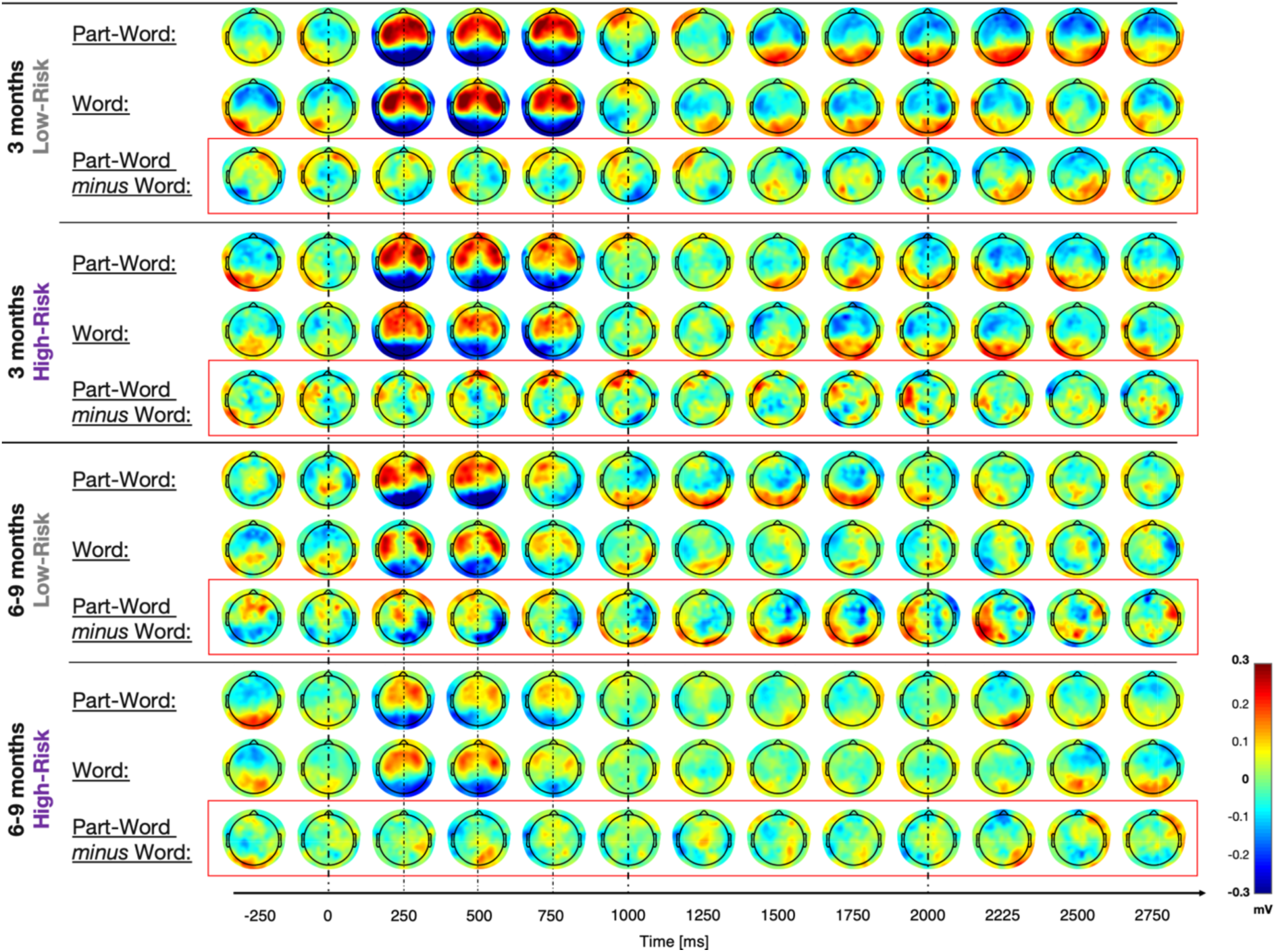
ERP topographies in each group at 3mo and 6-9mo.

**Figure S8:**
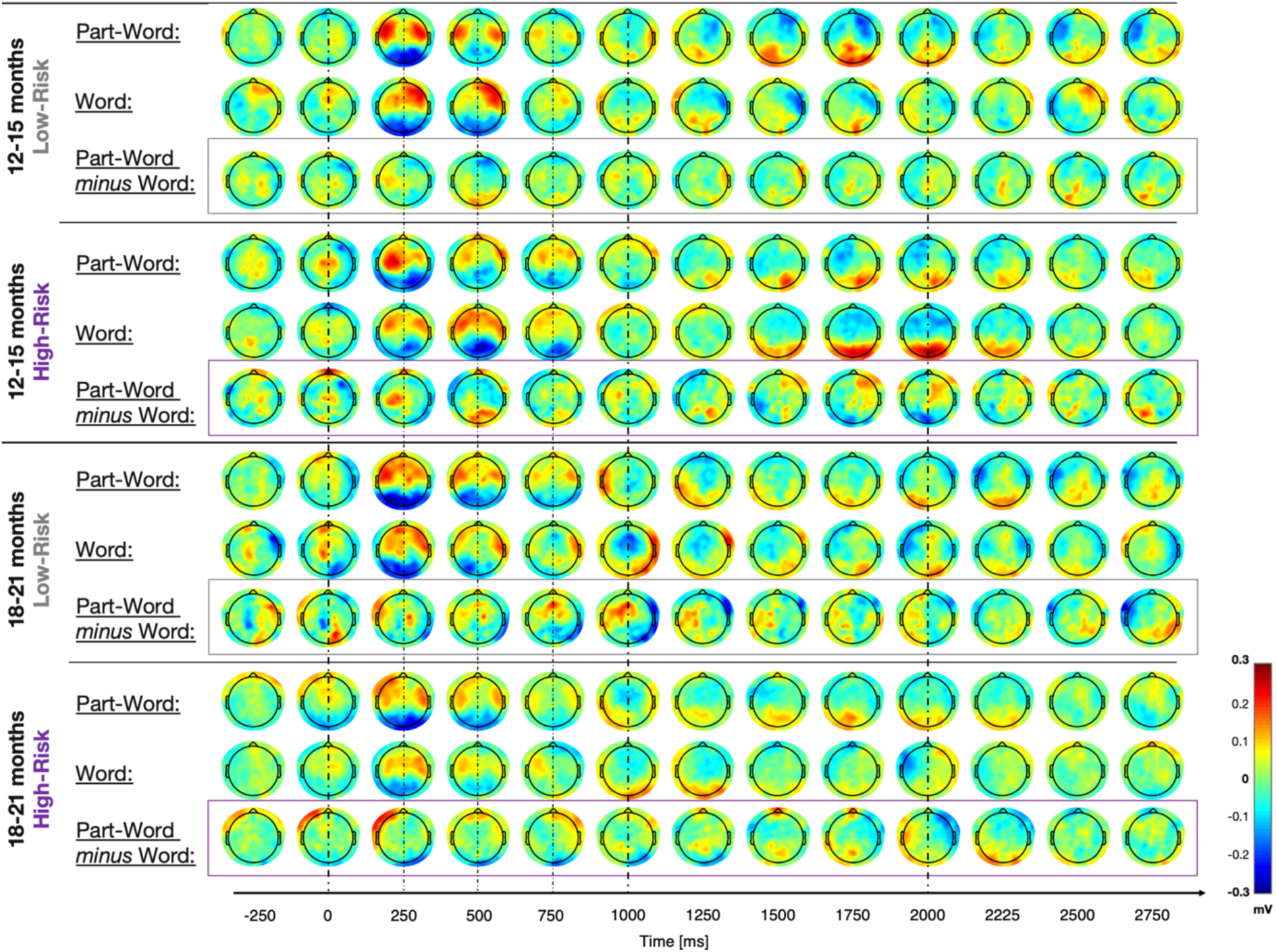
ERP topographies in each group at 12-15mo and 18-21mo.

